# Evidence for two main domestication trajectories in *Saccharomyces cerevisiae* linked to distinct bread-making processes

**DOI:** 10.1101/2020.05.28.120584

**Authors:** Frédéric Bigey, Diego Segond, Anne Friedrich, Stephane Guezenec, Aurélie Bourgais, Lucie Huyghe, Nicolas Agier, Thibault Nidelet, Delphine Sicard

## Abstract

Despite bread being one of the most historically and culturally important fermented products, its history and influence on the evolution of associated microbial species remains largely unknown. The first evidence of leavened bread dates to the second millenium BCE in Egypt and since, the art of bread-making developed and spread worldwide. Nowadays, leavened bread is made either by using a pure commercial culture of the yeast *Saccharomyces cerevisiae* or by propagating a sourdough, which is a mix of flour and water spontaneously fermented by yeast and bacteria. We studied the domestication of *S. cerevisiae* populations originating from industry and sourdough and tested whether these different bread-making processes led to population divergence. We found that the origin of *S. cerevisiae* bakery strains is polyphyletic with 67 % of strains clustering in two main clades: most commercial strains were tetraploid and clustered with strains having diverse origins, including beer. By contrast, most sourdough strains were diploids and found in a second clade of strains having mosaic genomes and diverse origins including fruits, or clinical and wild environments. When compared to the others, sourdough strains harboured in average a higher copy number of genes involved in maltose utilization, a common sugar produced from dough starch. Overall, a high level of gene flow from multiple contributors was detected. Phenotyping of bakery and non bakery strains further showed that sourdough and industrial bakery populations have undergone human selection for rapidly starting fermentations and for high CO_2_ production. Interestingly, sourdough strains also showed a better adaptation to a sourdough mimicking environment, suggesting that natural selection occurred as well. In summary, our results revealed that the domestication of bakery yeast populations has been accompanied by dispersion, hybridization and divergent selection through industrial and artisanal bakery processes. In addition, they unveiled for the first time a case of fungus domestication where species divergence occurred through autotetraploidisation.

## Introduction

The domestication of microbes is an ancient process that has accompanied fermented food processing since at least Neolithic times, when plant and animal domestication first occurred [1–7]. Until recently, the evolutionary history of domesticated microbes was poorly documented[2,3]. Most studies focused on the filamentous fungi *Aspergillus oryzae* used in rice and soya fermentation and *Penicillium roqueforti* used for making blue cheese [8–10] as well as on the *Saccharomyces cerevisiae* yeast model species. This last species is found in many natural habitats (soil, tree bark, water…) and has been domesticated for the production of a large diversity of fermented drinks (wine, beer, sake, cachaça, coffee, fermented milk) and foods (bread, cocoa, olives) [4,11]. Wild populations isolated from natural habitats present a broader and distinct genetic diversity than populations isolated from anthropogenic environments, suggesting that the latter were selected from the wild by humans for food processing [12,13]. The China/far East Asia area may likely be one center of origin of the domesticated populations [14].

The domestication of *S. cerevisiae* has been well described for wine, beer, sake, cachaça, cocoa, and coffee but surprisingly not for bread [14–16]. Domestication for making different products has led to a genetic diversification of strains that group together according to the fermentation type, but it also led to phenotypic divergence. Indeed, parallel domestication processes occurred for different beverages and foods although some gene flow between domesticated strains have been detected [12]. Wine strains and the closely related group of flor strains likely have a single origin [17–20]. Sake strains also appear to have evolved from a single origin [12,15,21] while cachaça strains evolved from wine strains through a secondary domestication process [22]. Coffee, cocoa and beer strains have a more complex evolutionary history where both migration and selection played major roles [23–27]. Several genetic signatures associated with human selection have been detected in all these domesticated populations, including SNPs [19,23,27], gene duplication [23,28,29], horizontal gene transfer [17–19,30], and genome hybridization [21]. Despite the cultural and historical importance of bread, the study of bakery strains domestication has been neglected. This might be related to the fact that several industrial bakery yeast starters have been found to be autotetraploid [31], which renders population genomic analysis complicated [32,33].

The earliest evidence of leavened bread was found during antiquity in the second millennium BCE in Egypt [34] and in the first millenium BCE in North West China [35]. Since then, the art of making leavened bread developed during ancient and medieval ages and was disseminated throughout the Mediterranean and in Middle East countries (Carbonetto et al. 2018). At that time, bread making consisted in mixing flour, water, and sourdough, a mix of flour and water containing fermenting microbes. In the 19th century, the industrialization of food production and the advent of microbiology as a science resulted in the production of pure yeast cultures that were used as starter to make bread. The production of bread made with commercial *S. cerevisiae* yeast starter soon spread all over the world. Yet, nowadays, global changes and the increase in the frequency of non-communicable diseases related to modern diets (including type 2 diabetes, obesity, food allergies) have led to a renewed interest in traditional methods of bread making and in its local production. The appeal of traditionally prepared breads is underpinned by research showing improved flavour and nutritional benefits in sourdough bread made by artisanal bakers [36–38]. Therefore, both ways of making leavened bread are currently found and bakers either use commercial yeasts, or natural sourdoughs.

Natural sourdough is made from flour and water and maintained by recurrent addition of flour and water, a process called backslopping. Sourdough contains a microbial community consisting of lactic acid bacteria and yeasts with a ratio of 100:1 on average [39]. One or two prevailing species of lactic acid bacteria and one prevailing yeast species are usually found. The yeast species found in sourdough mainly belong to the genera *Saccharomyces sensus stricto, Kazachstania, Pichia, Torulaspora*. Worldwide, *S. cerevisiae* is the most widespread species found in sourdoughs [40] made by bakers, but also in sourdough made by farmers-bakers [41,42]. It can be found as the dominant species in sourdough, even in bakeries where no industrial starter is used, suggesting that the species may colonize a sourdough from the bakery’s environment or the baker’s hands [43]. Therefore, bakery strains of *S. cerevisiae* may have undergone different domestication processes. While industrial bread production may have led to the breeding and selection of homogeneous lineages of yeast starters, artisanal bread making may have selected strains through the continuous, long-term maintenance of sourdough microbial communities. These two types of domestication processes also apply to many different fermented foods. [2]. Until recently however, study of fungi domestication mostly revealed footprints of industrial selection [9]. The renewed interest in traditional sourdough bread production makes the *S. cerevisiae* bakery populations an excellent model for the study of the impact of both artisanal and industrial practices on fungi evolution and adaptation.

The objective of this study was to investigate the evolutionary history of *S. cerevisiae* isolated in bakeries. We examined whether bakery strains had either a single or several genetic origins and studied the genetic diversity and genetic relationship of commercial and natural sourdough strains. In addition, we studied to what extent gene and genome duplication have been involved in the domestication of *S. cerevisiae* in bakeries. We found that bakery strains are polyphyletic and found in clades that also contain strains from other domesticated environments, suggesting no specific origin for bakery strains. Except for a few strains that clustered with wine or African beer fermentation strains, bakery strains were mostly grouped in two main clades, each composed of two subgroups. One mostly grouped commercial strains while the other mostly contained sourdough strains, suggesting different domestication roads for commercial and sourdough strains. Commercial strains appeared to be most often tetraploid and to display a shorter fermentation latency phase while sourdough strains appeared to have most often duplication of Maltose and Isomaltose maltase and permease genes, revealing different genetic and phenotypic signatures for industrial and artisanal selection. A overall high proportion of admixture was detected and some sourdough strains clustered together with commercial strains, suggesting that gene flow is also an important process in the evolution of bakery strains.

## Results

### Prevalent tetraploidy and aneuploidy in commercial bakery strains

Polyploidization, which refers to the multiplication of a complete chromosome set, has been found to be associated with domestication in plants, but also in *Saccharomyces cerevisiae* beer strains [23,44]. A previous analysis of 26 *S. cerevisiae* strains originating from diverse fermented products showed that most bakery strains analyzed were autotetraploid, suggesting that bakery yeast domestication was also associated with polyploidization [31]. We therefore analysed the ploidy of a set of 229 bakery *S. cerevisiae* strains (**Table S1**) using a combination of microsatellite typing and flow cytometry analysis. Thirty-one strains were commercial yeasts and 198 were isolated from European sourdoughs collected in Italy, Belgium and France. An overall high level of tetraploidy (40%) and aneuploidy (17%) was observed (**Table 1**). We found that commercial strains were significantly (two sided Fisher’s exact test, p < 0.001) more frequently tetraploid (68%) than sourdough strains (35%). On the other hand, we did not observe any significant difference in ploidy distribution between the 198 sourdough strains isolated in Belgium, France and Italy (two sided Fisher’s exact test, P > 0.05, **Table 2**).

**Table 1:** Ploidy variation in studied strains as revealed by fow cytometry and microsatellite typing

| origin | ploidy |  |  | total |
| --- | --- | --- | --- | --- |
|  | 2n | 4n | aneuploid |  |
| commercial | 5 | 21 | 5 | 31 |
| sourdough | 94 | 70 | 34 | 198 |
| total | 99 | 91 | 39 | 229 |

**Table 2:** Ploidy variation in sourdough strains isolated from France, Italy and Belgium, as revealed by both flow cytometry and microsatellite typing

| origin | ploidy |  |  | total |
| --- | --- | --- | --- | --- |
|  | 2n | 4n | aneuploid |  |
| France | 66 | 37 | 26 | 129 |
| Italy | 18 | 24 | 7 | 49 |
| Belgium | 10 | 9 | 1 | 20 |
| total | 94 | 70 | 34 | 198 |

To study whether tetraploidy promoted the adaptation of *S. cerevisiae* to a bakery environment, competition experiments between tetraploid and diploid strains were carried out in synthetic sourdough media. Commercial and sourdough strains of each ploidy level (**Table S2**) were included in the analysis to test not only the effect of ploidy but also the effect of commercial/sourdough origin. No evidence of fitness gain for tetraploids was found (t_43_=-1,4288, P=0.16, **Figure 1**). To test whether tetraploids provide a benefit for bakers’ practices, the effect of ploidy on fermentation kinetics was then analyzed. There was no significant effect of ploidy level on the maximum cumulative CO_2_ production released at the end of fermentation (CO_2_max, F_1,68_=0.1, P=0.749), maximum CO_2_ production rate (Vmax, F_1,68_=0.62, P=0.434) and time at Vmax (tVmax, F_1,68_=0.067, P=0.797) parameters (**Figure 2A,C,D**). By contrast, there was a significant effect of ploidy on the latency phase of CO_2_ production (time necessary to release 1g of CO_2_; F_1,68_= 7.01, P=0.01), while no significant effect of the origin of the strain (sourdough/commercial) was found (F_1,68_= 3.48, P=0.07). On average, tetraploids started fermentation earlier than diploids (**Figure 2B**).

**Figure 1:**
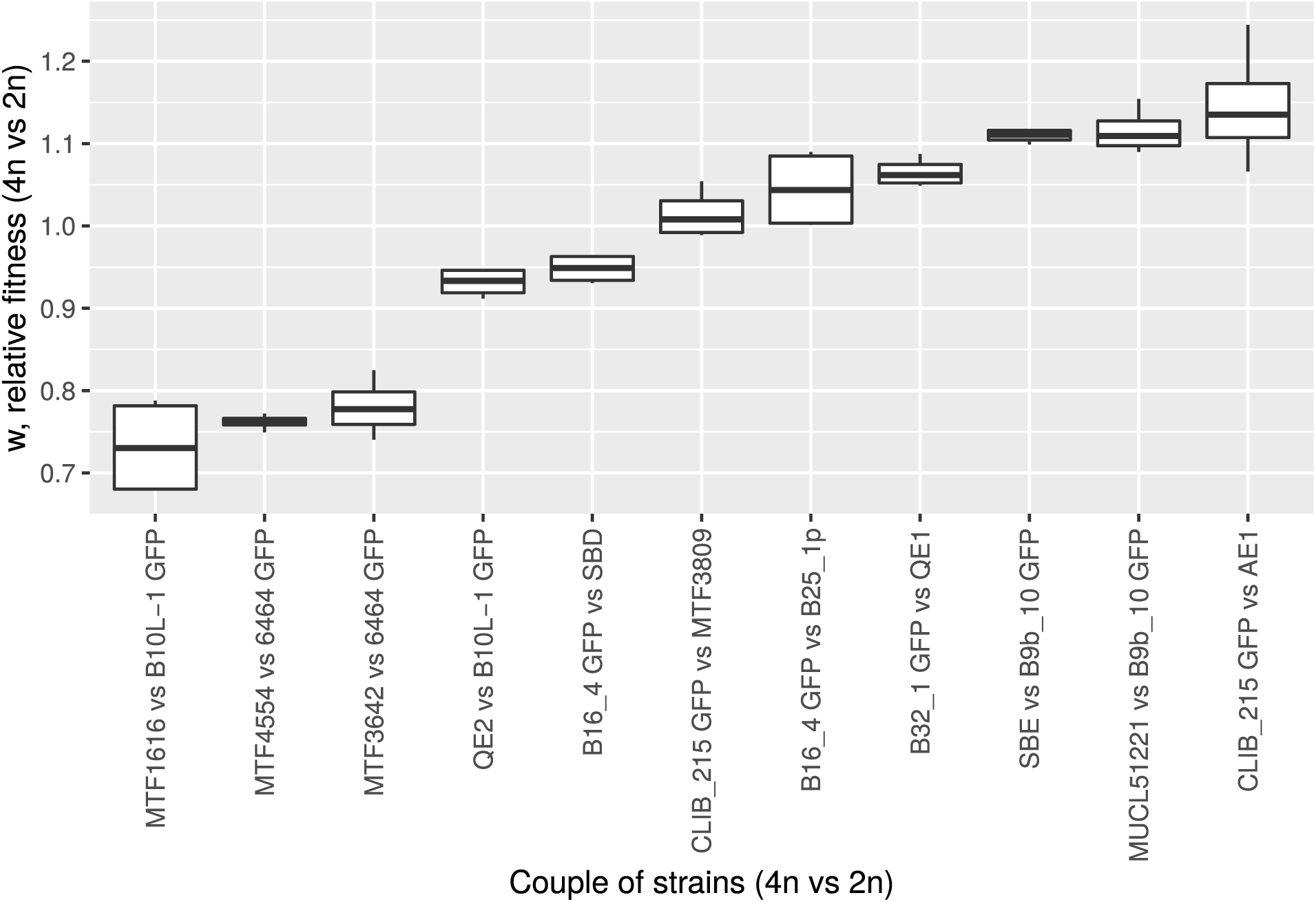
Effect of ploidy on fitness. A diploid and a tetraploid strains were cultivated in competition and relative fitness of 4N over 2N was computed after 24 h of fermentation in synthetic sourdough medium.

**Figure 2:**
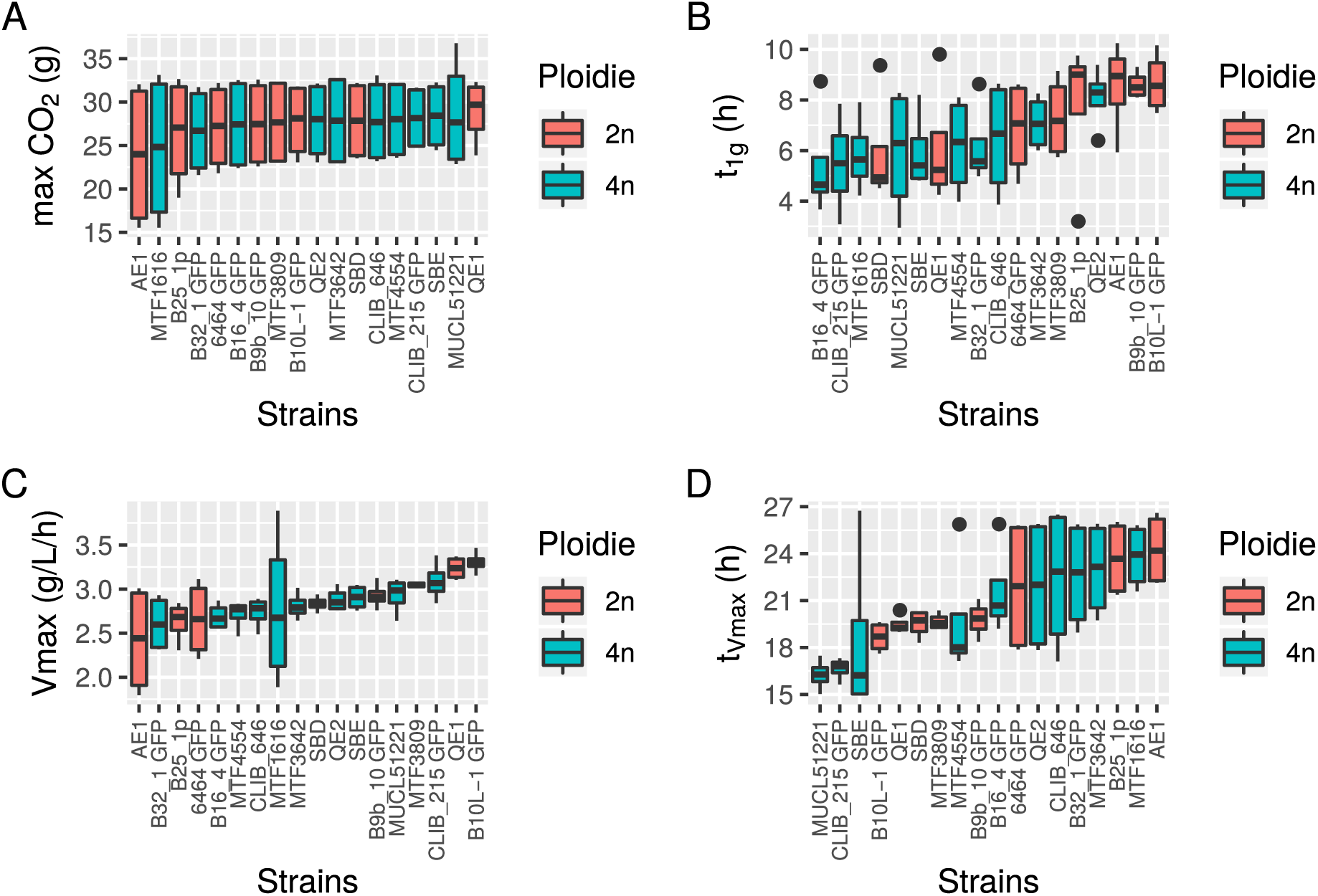
Effect of ploidy on fermentation kinetics. Diploids or tetraploids were cultivated in synthetic sourdough medium and CO_2_ released was monitored by weight loss. Four parameters were then estimated: **A**, maximum CO_2_ release (g); **B**, fermentation latency-phase time (h) (time elapsed between inoculation and the beginning of the fermentation calculated as 1g of CO_2_ release); **C**, the maximum CO_2_ production rate Vmax (g/L/h) and **D**, the time of the maximum CO_2_ production rate (h).

### Bakery strains are polyphyletic and present admixture

The evolutionary history of bakery strains was first studied on diploids using genomics. We examined the genomes of 68 bakery *S. cerevisiae* strains that included 17 newly sequenced diploid sourdough strains and a representative set of 51 previously sequenced bakery strains (**Table S5**). We studied the population structure of bakery strains based on 33,032 biallelic SNPs using fastStructure (**Figure 3**). This analysis yielded a most-likely population structure with 6 groups (likelihood -0.756). Groups P3 and P4 both contained sourdough strains, with a minority of other strains however for group P4. On the other hand, groups P2 and P6 were mostly composed of commercial strains. Finally, groups P1 and P5 comprised both sourdough and commercial strains. This genetic structuration was also observed using DAPC, which do not rely on any life-history traits and evolutionary assumptions (**Figure S1**) and on a maximum-likelihood phylogenetic tree (**Figure 4**), except for five strains that clustered on one side of the tree, outside of their group defined by fastStructure. Overall, there was no clustering according to the country of origin of the strains.

**Figure 3:**
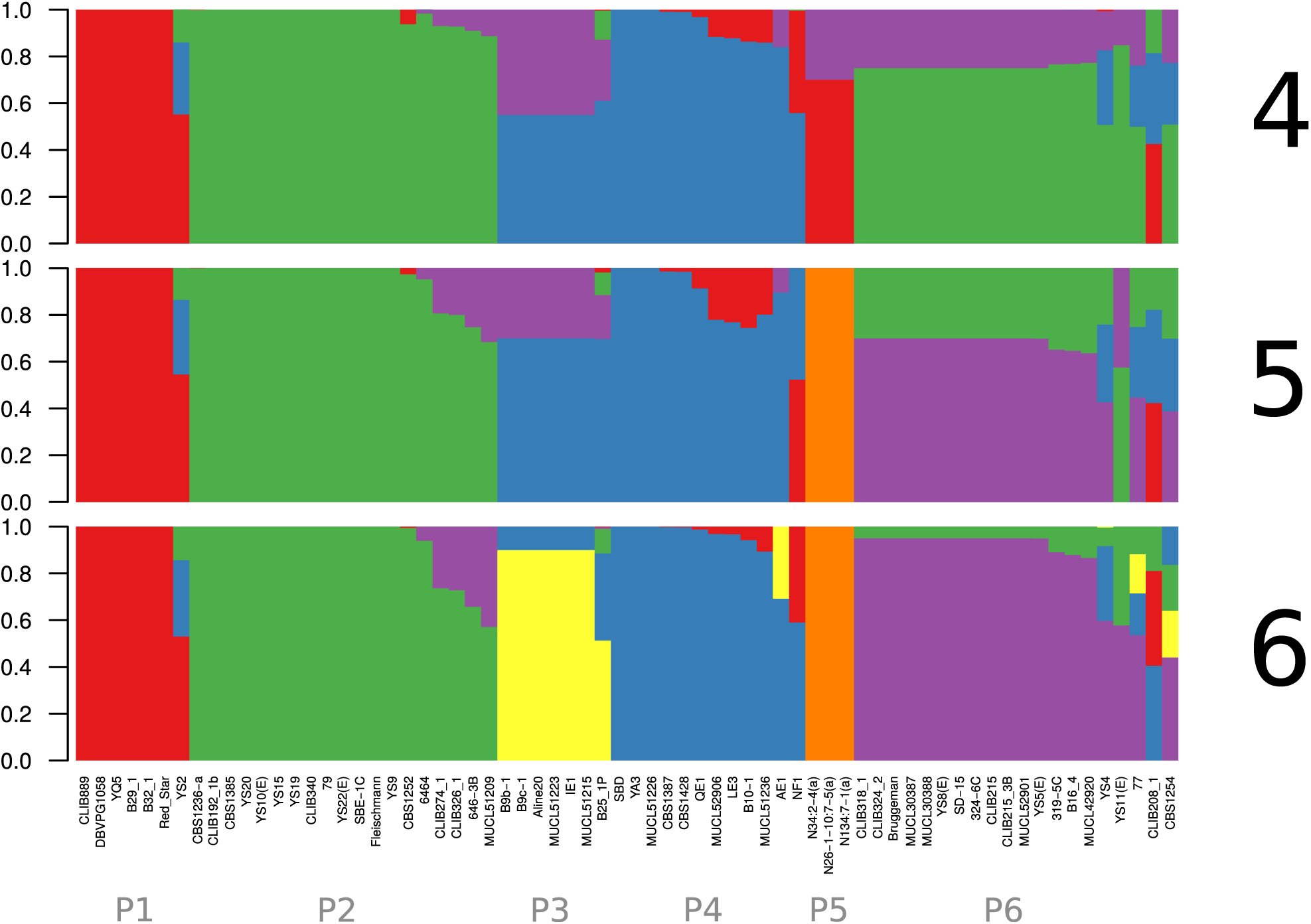
Population structure obtained from 33,032 biallelic SNPs from 68 bakery strains using fastStructure. The vertical axis depicts the fractional representation of resolved populations (colors) within each strain (horizontal axis) for K assumed ancestral populations. K = 6 maximizes the marginal likelihood (−0.756) and best explains the structure.

**Figure 4:**
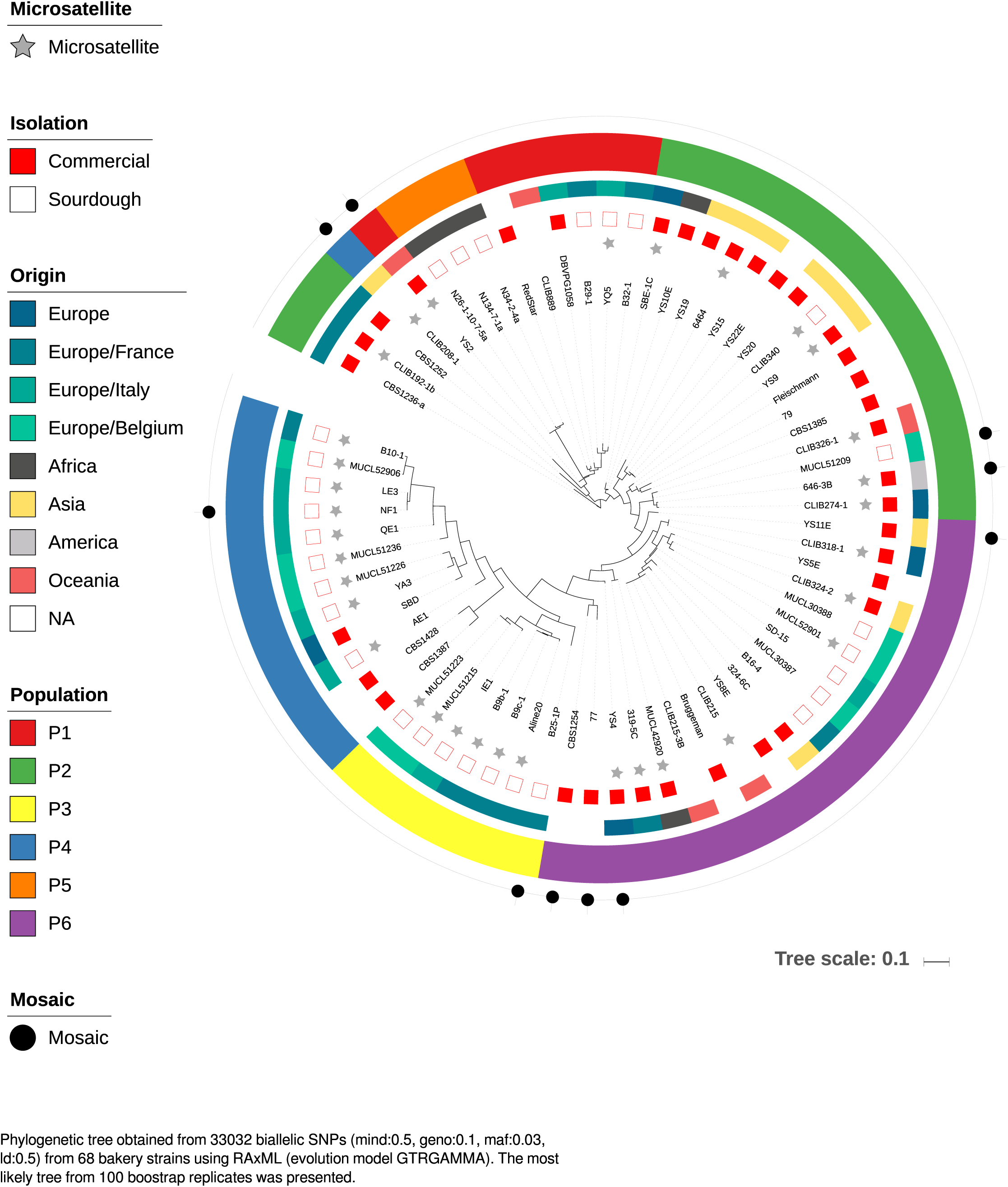
Maximum likelihood phylogenetic tree obtained from 33,032 biallelic SNPs from 68 bakery strains using RAxML (evolution model: GTRGAMMA). The most likely tree from 100 bootstrap replicates is presented. Groups P1-6 are defined in **Figure 1**.

We then analyzed the genetic relationship of bakery strains with the previously analysed 1,011 worldwide collection of *S. cerevisiae* **[12]**. Adding extra bakery genomes to the 1,011 genomes tree did not change the clustering defined in **[12]** (**Figure 5**). The bakery strains/genomes were distributed between the “Wine/European”, “African beer”, “Mixed origin”, “Mosaic region 3”, “Asian fermentation”, and “Mosaic region 1” clades. The P1 bakery group defined by fastStructure (7 bakery strains) was included in the “Wine/European” clade, while group P5 (3 bakery strains) was found in the “African beer” clade. The bakery groups P2 and P6 (23 strains, 7 from sourdough) were both located within the “Mixed origin” clade that also included beer strains, clinical strains, and strains isolated from water, fruits, tree leaves and natural environment. These two groups are indeed closely related and were not distinguished by fastStructure when the number of assumed ancestral groups K equalled 4 (**Figure 3**). Finally, the bakery groups P3 and P4 (19 strains, 16 from sourdough) were located within a group of mosaic strains that includes strains isolated from wine, sake, insect, palm wine, fruit, or clinical, fermentation, distillery, natural environment. These two groups are also closely related and were not distinguished when K equalled 5 (**Figure 3**).

**Figure 5:**
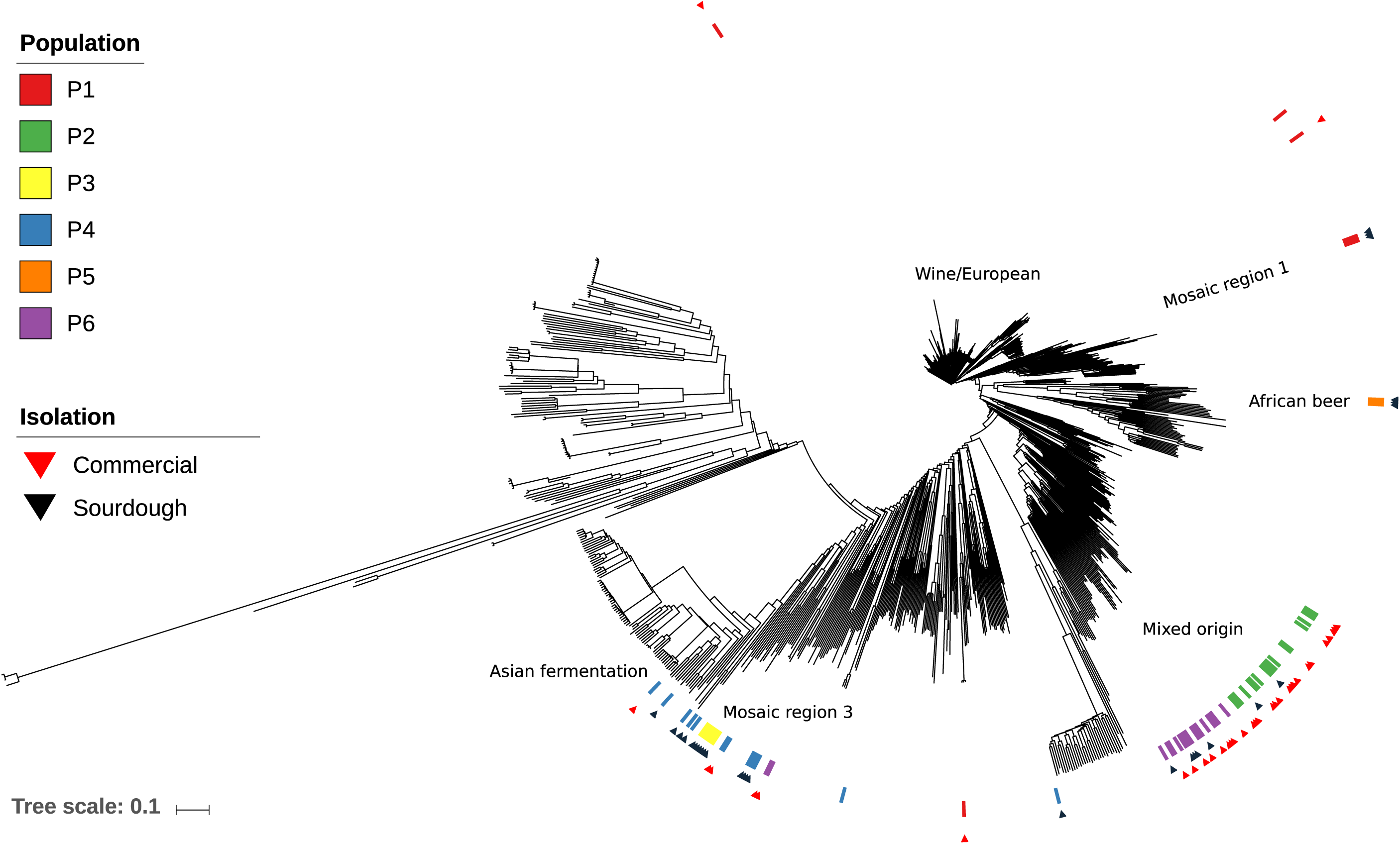
Phylogenetic tree obtained from SNPs from strains of the *S. cerevisiae* 1,011-genomes project (Peter et al. 2018). 17 newly sequenced diploid sourdough strains genomes and a representative set of 51 previously sequenced bakery strains extended the dataset to a total of 68 bakery yeast genomes (**Table S5**). Groups P1-6 are the same as in **Figure 1**. The names of the clades are taken from the 1,011-genomes project as described in **[12]**.

To analyse the degree of admixture of bakery strains, we ran Admixture on the bakery genomes as well as on 90 genomes chosen across the 1,011 genomes tree clades [12]. The strains were chosen among all the clades except those not found to contribute to the bakery-strains containing clades (**Figure 6, Table S6**). A total of 48 bakery strains out of 68 presented some level of admixture varying between strains, and reaching up 70% of the genome. For a single strain, from 2 to 11 ancestral populations were admixed. There was evidence of admixture between bakery strains of “Mosaic region 3” and “Mixed origin” clades. In addition, there was evidence of admixture with other clades and contributors were found to belong to the genetic clades “Asian fermentation”, “Wine/European”, “Ale beer”, “Brazilian bioethanol”, “Mosaic beer” and “African beer”. One contributor however could not be identified, suggesting the presence of an extinct or otherwise uncharacterized *S. cerevisiae* population. Novel alleles derived from unknown or extinct populations were also found in ale beer strains (Fay et al. 2019). The high degree of admixture suggested that dispersion and hybridization were parts of the main drivers of bakery strains evolution.

**Figure 6:**
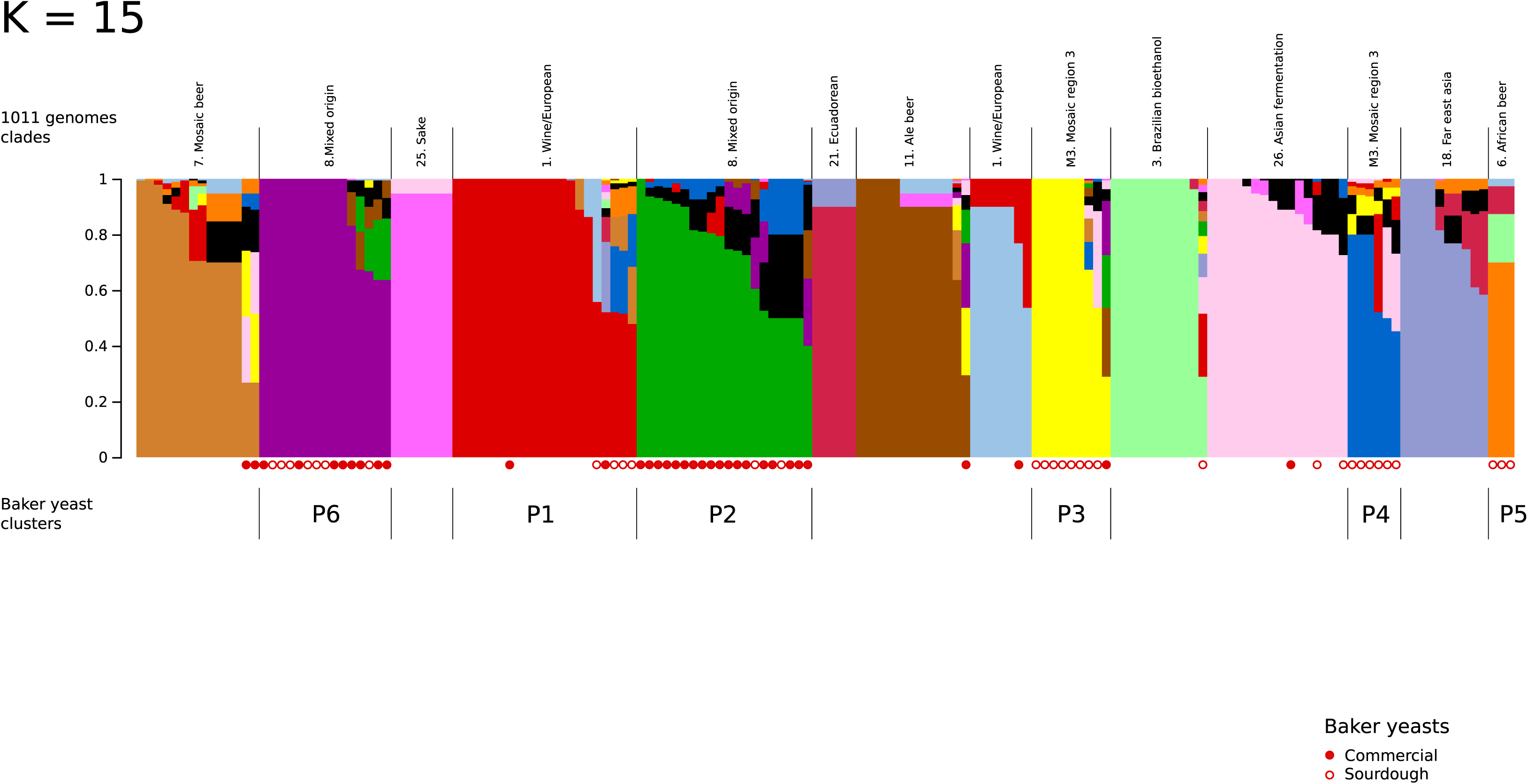
Population structure obtained from 48,482 biallelic SNPs from 157 genomes (including 68 bakery strains; **Table S6**) using Admixture software. The vertical axis depicts the fractional representation of resolved populations (colors) within each strain (horizontal axis) for K assumed ancestral populations. The value of K = 17 exhibited the lowest cross-validation error compared to other K values and best explained the structure. Groups P1-6 are defined in **Figure 1**. The clades from the 1,011-genome project are those described in [12].

The bakery strains of the “Mixed origin” and “Mosaic region 3” clades differed by the origin of some introgressions. Introgression from the “Ale beer” clade was only detected in commercial and sourdough strains of the “Mixed origin” clade (28% of the “Mixed origin” bakery strains; **Figure 6**). Introgression from the “Asian fermentation” population was only found in bakery strains of the “Mosaic region 3” clade (60% of the “Mosaic region 3” bakery strains; **Figure 6**). This last result was confirmed by an analysis with Treemix **[45]** that evidenced a gene flow between “Asian fermentation” and “Mosaic region 3” bakery populations (w = 38%, three-population f3 test, Z = -27; **Figure S2**).

### An increased copy number of maltase and isomaltase, transporter and regulator genes in the sourdough strains clade

Large copy number variations were previously detected in *S. cerevisiae* [12]. We analysed large CNV on our bakery strains (**Table S7**) and found that twenty-six strains (40%) displayed major chromosomal rearrangements or aneuploidies. Chromosome 9 was the most affected by CNV (10 strains out of 26). In addition, we analyzed the copy number of genes involved in maltose, iso-maltose and sucrose assimilation. These carbohydrates are common in cereal products which may have led to the selection of an increased number of genes involved in their assimilation in bakery strains. Gene copy number was compared between bakery and non-bakery strains within each clade containing bakery strains to eliminate the genetic structuration bias (**Figure S3**). Analysis was first performed on the MAL maltose gene cluster [46,47]. In *S. cerevisiae*, the maltose gene cluster is composed of three genes, encoding the maltose transporter (permease, MAL1), maltase (MAL2) and transcription regulator (MAL3). The genes involved in maltose utilization are represented in five well-described MAL loci located on subtelomeric regions [46,47]. The presence of just one MAL locus is sufficient to allow for maltose fermentation [48,49]. In the “Mosaic region 3” clade, where most bakery strains were isolated from sourdough, the number of copies of the *MAL12* (Wilcoxon rank sum test, p-value < 10^−3^) and *MAL32* (p-value < 10^−4^) maltase genes was on average significantly higher in bakery strains than in non-bakery strains (**Figure S3A,B**). The same was observed for the maltose permease gene *MAL31* (p-value < 10^−4^). The same analysis was then performed for the isomaltase genes (**Figure S3C**). A significant increase in copy number for *IMA1* (p-value < 10^−6^), *IMA3* (p-value < 10^−4^) and *IMA4* (p-value < 10^−4^) in bakery strains compared to non-bakery strains was also observed in the “Mosaic region 3” clade. In the “Mixed origin” clade where many commercial bakery strains were located, we did not detect any increase in of copy number for genes in the maltose and isomaltose gene clusters in bakery strains compared to others. These results suggest selection for a better assimilation of maltose and isomaltose in sourdough, where starch degradation release maltose. In the same way, adaptation to beer environment was found to be associated to an increased copy number of the maltase gene cluster **[50]**.

### Phenotypic signatures of domestication

To test further whether sourdough strains have undergone human selection, we compared fermentation performance between bakery and non-bakery strains and analyzed phenotypic convergence among bakery strains. Fourteen sourdough strains, six commercial strains and six non-bakery strains of diverse origins and genetic groups were fermented in a synthetic sourdough medium (**Table S8**). As previously, four fermentation parameters relevant to bread-making were studied: time necessary to release 1g of CO_2,_ the maximum cumulative CO_2_ production released at the end of fermentation (CO_2_max), maximum CO_2_ production rate (Vmax) and time at Vmax (tVmax). In addition, the number of cells after 27h of fermentation was used as proxy for absolute fitness. We found that non-bakery strains had significantly lower maximum cumulative CO_2_ production, lower maximum CO_2_ production rate and started fermentation later than sourdough and commercial bakery strains (**Table S9, Figure 7**) showing that both sourdough and commercial strains have been selected for better performance in fermentation. Commercial bakery strains performed better than sourdough strains in terms of fermentation onset and CO_2_ production. However, sourdough strains had a maximum CO_2_ production rate as high as commercial strains. Moreover, sourdough strains reached a significantly higher population size than commercial strains suggesting they display increased fitness in a synthetic sourdough medium compared to industrial yeasts (**Figure S4**). Overall, these results confirmed that sourdough strains were domesticated by artisanal and farmer-bakers and are better adapted to their environment than other strains.

**Figure 7:**
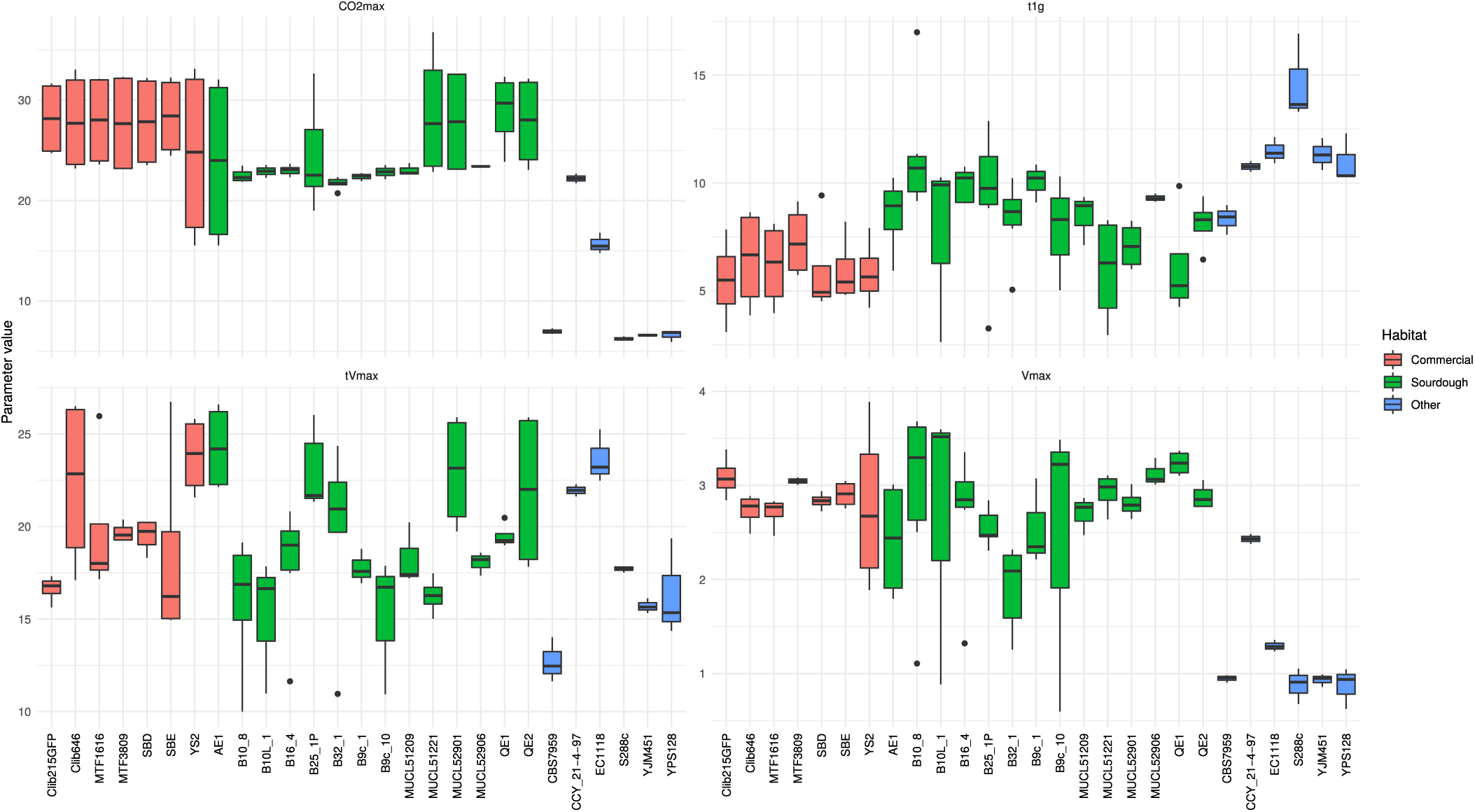
Phenotypic variation of fermentation kinetics in a sourdough synthetic medium among sourdough, commercial and non-bakery strains (other). Four parameters were estimated: the maximum CO_2_ release (g), the fermentation latency-phase time (h) (time between inoculation and the beginning of the fermentation calculated as 1g of CO_2_ release), the maximum CO_2_ production rate Vmax (g/L/h) and the time of the maximum CO_2_ production rate.

### Genetic relationships between diploids and tetraploids

To analyse the genetic relationship between diploids and tetraploids, we studied genetic diversity in 229 bakery strains (31 commercial and 198 isolated from sourdough; **Table S1**) with 15 microsatellite markers. A total of 31 strains were analyzed using both microsatellites and genomic sequences, which allowed the comparison of genomic and microsatellite data. The microsatellite genetic relatedness tree based on the Ritland relatedness coefficient [51] showed a clustering concordant with the genomic groups (**Figure 8**). Commercial strains clustered in groups P2 and P6 while sourdough strains were scattered all over the tree. Groups P3 and P4, mostly represented by sourdough strains, contained mostly diploids (76 diploids strains and 5 tetraploids strains) while groups P2 and P6 contained mostly tetraploids. Genotypes were clustered according to the sourdough from which they have been isolated. Strains from the same sourdough were either all diploids (sourdoughs B9b, Al) or all tetraploids (sourdough F). Interestingly two sourdoughs (B9c, B10L) contained both diploids and tetraploids. In one case, there was one tetraploid and seven diploids. In the other case, there was four diploids and 9 tetraploids. The diploids and tetraploids were very closely related suggesting they derived from each other rather than from different introduction events.

**Figure 8:**
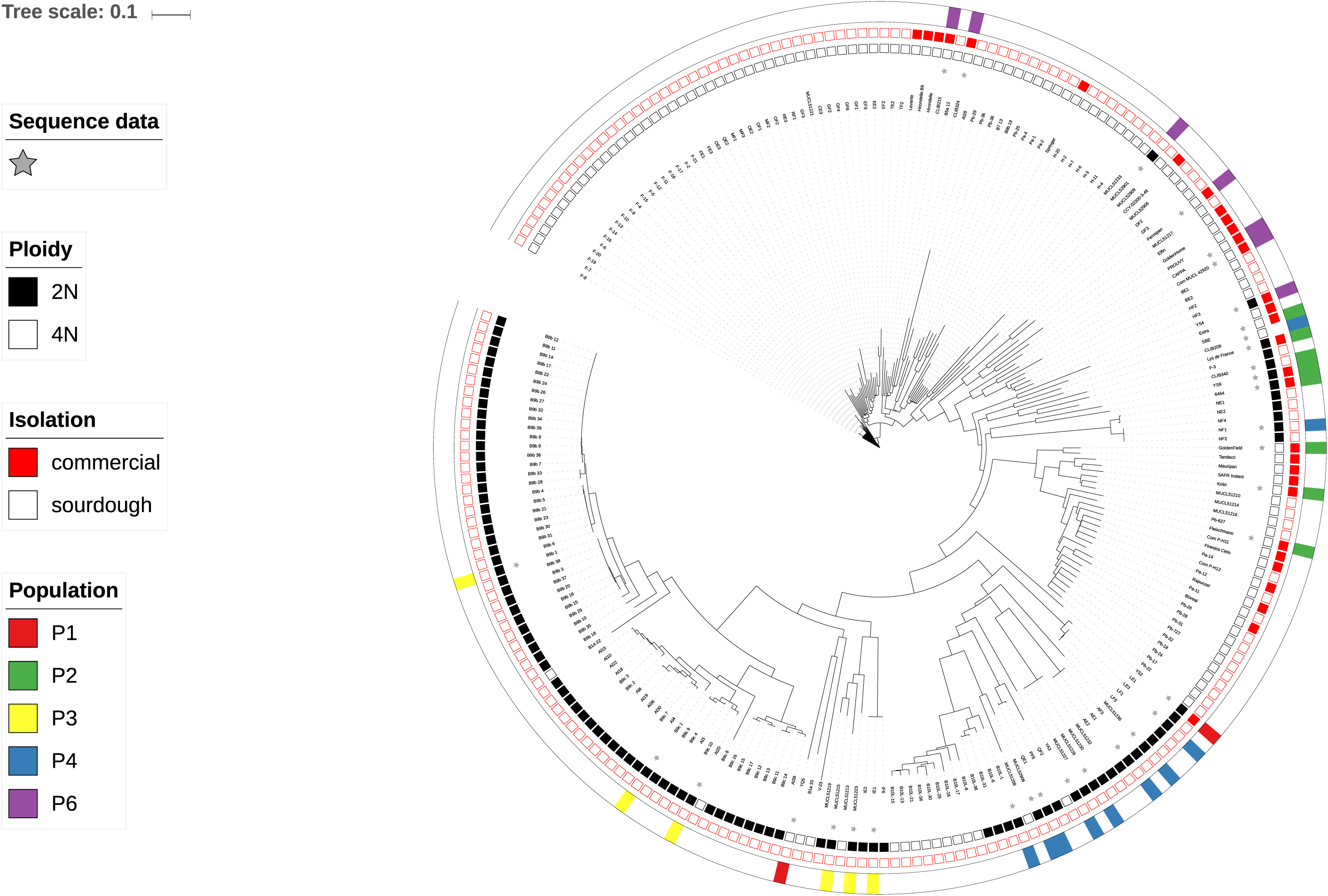
NJ tree computed using the Ritland coefficient of relatedness obtained from microsatellite typing of the 229 bakery strains. Groups P1-6 are defined in **Figure 1**. Grey stars indicate strains for which a genome sequence was obtained.

Further analysis of population structure using an Analysis of Molecular Variance (AMOVA) [52] allowed the joined analysis of diploids and tetrapoids [53]. First, we studied the differences between bakery origin (commercial *vs* sourdough strains). Only 11% of the variation was found between bakery origins (**Table S10**) while the variation within bakery origin was 89%. However, there was a significant structuration by bakery origin (permutation test p-value < 10^−3^), confirming the genomic and microsatellite clustering of most sourdough strains on one side and most commercial strains on the other. Then we focused on sourdough strains isolated in France, Belgium and Italy and examined whether there was any geographical structuration of sourdough strains according to their country of origin. To avoid unequal sampling between country, three strains per sourdough were randomly sampled. Most of the variance arose within each country (89%). However, a permutation test revealed significant differentiation between countries (permutation test p-value < 10^−3^). Significant differentiation between countries and similar distribution of variation were also found when all analyzed strains from France, Italy and Belgium were included. Finally, the distribution of genetic diversity within and between sourdoughs was analyzed for France, where a large number of yeast strains per sourdough were available. Most of the observed variance occurred between sourdoughs (76%), showing that genetic variation between sourdoughs exceeded genetic variation within sourdough. Permutation tests revealed a significant structuration according to the sourdough of origin (simulated p-value < 10^−3^). However, some intra-genetic sourdough diversity was also found as for example in sourdough B9b and Al, where a strain (B9b, Al28) clustered far from most of the strains of the sourdough revealing a dispersion event.

## Discussion

We report here the first broad analysis of bakery yeast domestication using a collection of 229 *Saccharomyces cerevisiae* bakery strains collected worldwide from industry or natural sourdoughs. We found that the origin of bakery strains is polyphyletic. Most bakery strains clustered in two main clades, suggesting that bakery strains have undergone at least two main domestication trajectories: one domestication trajectory appeared to have led to most commercial strains, while the other led to sourdough strains.

The domestication of commercial bakery strains was associated with at least one tetraploidization event and the selection for a shorter latency phase of fermentation. To our knowledge, this is the first time that selection and dispersion of widespread autotetraploids are associated with the domestication of fungi. Beer strains of *S. cerevisiae* were found to have a high rate of ploidy variation associated in addition to aneuploidy, but there was no evidence of worldwide spread of autotetraploids [12,50]. A change in chromosomal copy number is often observed in yeast whenever they adapt to new, stressful environments [44,54,55]. However, several studies showed genome instability in yeast polyploids. Chromosome loss, aneuploid mis-segregation event, chromosome translocation, and large chromosome rearrangement all occurred during the evolution of yeast polyploids [44]. Experimental evolution results showed that tetraploid ancestors converged into diploids in 1800 mitotic generations [56,57]. Here we found that the vast majority of commercial yeasts are autotetraploid as are also some strains isolated from sourdough. Bakery autotetraploids are in reproductive isolation with diploids and thus represent a new species (Albertin et al. 2009). The high proportion of tetraploids among commercial strains as well as the association of tetraploids with shorter fermentation latency phase suggest that deliberate artificial selection by industrial yeast geneticists could be at the origin of tetraploid bakery strains and that worldwide industrial distribution of these selected strains might have disseminated tetraploids in bakery environments. The high level of aneuploidy observed in bakery strains could result from the known instability of tetraploid genomes.

The domestication of sourdough strains was associated with an increase in copy number of the maltose and isomaltose transporter (permease) and maltase, isomaltase transcription regulator encoding genes. While CNVs are generally deleterious, they also appear to be a key mechanism that can enable adaptation during an episode of stringent selection. CNVs are widespread in domesticated plant and animal species [58] as well as in domesticated yeast populations (Legras and Sicard, 2011; Steensels et al. 2018). Because maltose and isomaltose are released through amylolytic starch breakdown, they represent an important carbon source in dough and may directly be linked to fermentation performance (duration and CO_2_ production). Therefore, natural selection in sourdough as well as unintentional selection by bakers for increased fermentation rate may both have selected for strains having an increased copy number of genes related to maltose and isomaltose utilization. A slight but significant geographic structuration of the genetic diversity was found according to sourdough and its country of origin suggesting an effect of bakery practices and environments.

Some sourdough strains were found in the commercial strains”Mixed origin” clade, and vice versa, suggesting that commercial starters may disseminate in natural sourdoughs. Some bakers may indeed add commercial starter to their sourdough or may contaminate their sourdough by using commercial yeast in the bakery house for making other bakery products such as pastries. Admixture between these two clades was also detected, suggesting gene flow occurrence between commercial and sourdough strains.

Bakery strains have been hypothesized to be genetically related to beer strains (Legras et al. 2007, Gallone et al. 2016). In the XVIIth century, baker’s yeast was reportedly provided by brewers **[4]**. Yet, although beer strains were found in seven clades out of the 26 clades structuring the 1,011 *S. cerevisiae* worldwide strain collection **[12]**, only two of these clades also contained bakery’s strains. First, the African beer clade contains sourdough strains isolated from maize dough coming from Ghana, suggesting that the same strains are used both to ferment maize dough and to brew beer. Second, the “Mixed origin” clade contains beer strains, commercial bakery strains from all over the world, a few sourdough strains from Belgium and France as well as strains from diverse other habitats (fruits, soil, water, humans, …). None of the bakery strains clustered with the “Ale beer” clades. However, some introgression from the Ale beer clade was detected in ten out of the 36 bakery strains of the “Mixed origin” clade indicating the presence of some gene flow between “Ale beer” and bakery strains. This clade mostly contained commercial strains suggesting that some Ale beer strains have been used as progenitors in the industrial breeding of bakery strains.

A previous study also proposed that bakery strains could originate from a tetraploidization event between ale beer and wine strains [16]. Moreover, a recent study found that ale beer strains were derived from admixture between populations closely related to European grape wine strains and Asian rice wine strains [59]. Here, all bakery strains but three clustered separately from the well-defined wine and sake lineages, clearly suggesting a distinct evolutionary history. The admixture analysis provided no evidence of gene flow between bakery and sake strains. However, there was evidence of small introgression from wine populations and from “Asian fermentation” populations. Interestingly, sign of introgression from wine populations were found both in sourdough and commercial strains while introgressions from “Asian fermentation” populations were detected only in sourdough strains (Figure 4). Artisanal bakers often experiment other fermentation processes, among which “Asian fermentation”, which may explain this finding.

Sourdough is a human-made habitat. One may consider that any yeast population originating from sourdough is domesticated since this environment would not exist without human intervention. Alternatively, one may consider that sourdough yeast populations can only be considered domesticated if some human selection has indeed occurred. Here, we provided evidence that sourdough strains are not only present in their environment by chance or by recurrent introduction but have been selected for better fermentation performance. They compared to commercial strains in terms of maximum CO_2_ production rate but reached a higher population size at the end of fermentation. These results suggested that sourdough strains are better adapted to a sourdough environment and provided interesting genetic resources for improving sourdough bread making process.

In summary, our study revealed that bakery strains have undergone at least two main domestication trajectories that mobilized different genetic events (tetraploidization, CNV), and selection targets (shortened latency phase of fermentation in industry, adaptation to maltose utilization and sourdough environment in artisanal bakery). We also showed that dispersion and gene flow is an important driver of bakery strains evolution and that different sources of introgression have occurred in sourdough and commercial strains (respectively “Asian” and “Ale beer” sources). Overall, this is the first time that the analysis of fungal domestication revealed that artisanal and industrial domestication led to divergent populations. This demonstrates the need of conserving different fermentation practices to maintain microbial genetic diversity.

## Materials and Methods

### Yeast collection and cultivation procedure

The collection of strains analysed is presented on **Tables S1, S5, S8**. One hundred and twenty-nine strains were collected from sourdoughs in France as previously described [42,60]. Twenty sourdough strains from Belgium isolated by [61] were kindly provided by the Belgium MUCL collection. Finally, 47 sourdough strains isolated by [62] in Italy were kindly provided by Fabio Minervini through the University of Perugia yeast collection (Minervini et al. 2012) and two sourdough strains from Sicily were kindly provided by Jean-Luc Legras [19]. Thirty-one commercial strains were ordered from different international yeast collection or bought as starters.

Cultures were performed in 10 mL of liquid YE medium and were inoculated with yeast cells either from frozen stocks (stored at -80 °C) or by picking a single colony from a previous culture plate.

### Ploidy analysis by flow cytometry

Strain ploidy was analyzed by flow cytometry. Namely, 10^7^ yeast cells, recovered at the beginning of stationary phase, were fixed in 70% ethanol for 16 h at 4 °C. They were washed with 200 μL sodium citrate buffer (50 mM, pH 7) and then dispersed in 1 mL of the same buffer. 100 μL of cell suspension were transferred to a microtube and treated with 1 μL RNAse A (100 mg/mL) for 2 h at 37 °C. Labelling was performed by addition of 400 μL of a staining solution (50 μg/mL propidium iodide in citrate buffer), and incubating for 40 min at 20 °C in the dark. Cells were recovered by centrifugation and resuspended in 500 μL citrate buffer.

Flow cytometry analysis of 30,000 cells was carried out using the MACSQuant® Analyzer from Miltenyi Biotec GmbH (Germany) and the data analysis by the MACSQuant® software. A haploid strain (S288C) and a tetraploid strain (Levante; (W. Albertin et al. 2009)) were used as calibration controls.

### Fitness and fermentation analysis

#### GFP labelling

To study the impact of ploidy and strain origin (sourdough/commercial) on fitness, three diploid and three tetraploid strains were tagged with GFP. The pFA6a-TEF2Pr-eGFP-ADH1-NATMX4 plasmid [63] was used as a template to amplify a cassette containing the TEF2 promoter, eGFP, ADH1 terminator and NATMX4 conferring resistance to clonNat. The PCR fragment obtained with primers GFPNATMXtoHO-for (5’-GCTATTGAGTAAGTTCGATCCGTTTGGCGTCTTTTGGGGTGTAACGCCAAGATCTGTTTAGCTTGC CTTGTC-3’) and GFPNATMXtoHO-rev (5’-GAGGCCCGCGGACAGCATCAAACTGTAAGATTCCGCCACATTTTATACACTCATGAATTCGAGCTCGTTGTC-3) was inserted into the HO locus of the selected strains. All *S. cerevisiae* strains used here are listed in **Table S3**. The fitness and fermentation cost of carrying the GFP construction was assessed by competing the GFP-labelled and its unlabelled ancestral strain, as well as by comparing fermentation kinetics of these two strains. Relative fitness of the GFP-labelled strain relative to the unlabelled one was not significantly different from 1 (One-sample t-test, two-sided, **Table S4**) and neither did GFP labelling change fermentation kinetics parameters (ANOVA, **Table S4**).

#### Competition and Fermentation conditions

Competition between GFP-labelled and unlabeled strains and single strain fermentations kinetics were performed in a sourdough synthetic medium (SSM) that was adapted from [64] to better mimic the average composition of sourdoughs. The SSM contained, per liter: wheat peptone, 24 g; MgSO_4_.7H_2_0, 0.2 g; MnSO_4_.H_2_O, 0.05 g; KH_2_PO_4_, 4 g; K_2_HPO_4_, 4 g; Tween 80, 1 mL; glucose, 15 g; maltose, 35 g; cobalamine, 0.2 mg; folic acid, 0.2 mg; nicotinamide, 0.2 mg; pantothenic acid, 0.2 mg; pyridoxal-phosphate, 0.2 mg and thiamine, 0.2 mg. The pH was adjusted to 4.5 with citric acid and the solution was sterilized by steam pasteurization for 15 min. Sterile-filtered vitamin and sugar solutions were added after pasteurization. Overnight pre-cultures in YPD were titrated with a C6 flow cytometer (Accuri, BD Biosciences). For competition experiments, 5.10^5^ cells/mL of each strain were then inoculated in 15 mL SSM medium. For single strain analysis of fermentation kinetics, 10^6^ cells/mL of pre-culture were inoculated in 15 mL SSM media.

Fermentations were carried out at 24 °C with constant magnetic stirring (300 rpm) in 20 mL glass tubes closed with a filter tip to allow release of CO_2_. Fermentations were monitored during 24 h for CO_2_ release, by measuring weight loss every 40 min using an automated robotic system [65].

At the end of fermentation, cultures were centrifuged and pellets were resuspended in PBS for flow cytometry analysis (C6 cytometer, Accuri, BD Biosciences). Population size and cell viability were determined as described in [66]. Relative fitness of tetraploid vs diploid strains was estimated based on the proportion of GFP-labelled vs unlabelled strains in mixed cultures. GFP fluorescence (excitation 488 nm, emission 530 nm) was collected in the FL1 channel.

#### Statistical analysis

All competition and single strain fermentation were carried out with 3 or 4 replicates.

Relative fitness was calculated as

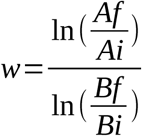

where w is fitness, A and B are the population sizes of the two competitors, while subscripts i and f indicate the initial and final time point (24h) of the competition kinetics [67]. To statistically compare the fitness of tetraploids and diploids, the mean relative fitness of tetraploids relative to diploids was compared to 1 using a one-sample t-test, two-sided (**Table S4**).

To test the effect of ploidy and strain origin (commercial/sourdough) on fermentation kinetics, the cumulative CO_2_ production curve was calculated and the kinetics of CO_2_ production rate over time was estimated. Four parameters were then estimated: the maximum CO_2_ release (g), the fermentation latency-phase time (h) (time between inoculation and the beginning of the fermentation calculated as 1g of CO_2_ release), the maximum CO_2_ production rate Vmax (g/L/h) and the time of the maximum CO_2_ production rate. The effect of ploidy and strain origin was tested for each kinetic parameter using the following linear model: *Y* _*ijk*_ =*μ* + *ploidy*_*i*_ +*origin*_*j*_+ *ploidy x origin*_*ij*_+*ε*_*ijk*_, where Y_ijk_ is the kinetics parameter variable, α_i_ is the fixed ploidy effect, β_j_ the fixed origin effect, γ_ij_ the interaction effect and ε_ijk_ the residual error.

To test the “habitat of origin” effect (sourdough/commercial/other) on fermentation kinetics, the same four kinetics parameters were estimated. The number of cells after 27h of fermentation was also analyzed as a proxy of absolute fitness. The following mixed linear model was used *Y* _*ijk*_ =*μ* + *Bloc*_*i*_+*Strain* _*j*_+*origin*_*k*_ +*ε* _*ijk*_, where Y_ijk_ is the fermentation kinetics or population size variable, Bloc_i_ is the random bloc effect, Strain_j_ the random strain effect, origin_k_ the habitat of origin fixed effect and ε_ijk_ the residual error. Tukey tests were used to compare pair of means.

### Genomic DNA extraction

Cells from an overnight culture were recovered by centrifugation of 5 mL of culture medium at 4500 rpm for 3 min. The yeast pellet was suspended in a sorbitol solution containing 20 μL Zymolyase 20T (1 mg/mL). Cell wall lysis was performed at 37 °C for 1 h. DNA extraction was carried out using the DNAeasy Blood & Tissue Kit (Qiagen).

### Microsatellite typing and analysis

In order to study the genetic diversity of diploid and tetraploid, 15 microsatellites loci previously described in [68] were used: C3, C4, C5, C6, C8, C9, C11, ScAAT1, ScAAT3, SCYOR267C, YKL172w, YKR072c, YLL049, YLR, YPL009c. Multiplexing was used in order to amplify 4 or 5 microsatellites per PCR run, with two distinct markings, one at 700 nm and the second at 800 nm. Once amplified, amplicons were diluted 1/20 in formamide, denatured for 5 minutes at 85 °C, then separated by electrophoresis on a 13 % polyacrylamide gel containing 39 % urea in 1.2X TBE buffer at 2000 V for 15 h (50 °C) on an automatic sequencer.

Analyses of microsatellite data were performed using Poppr version 2.8.2 [69]. A microsatellite genetic relatedness tree was constructed based on the Ritland relatedness coefficient [51]. Analysis of Molecular Variance (AMOVA) was used as a method of estimating population differentiation (Excoffier, Smouse, et Quattro 1992).

### Genome sequencing and read processing

Genome sequencing data were obtained for 68 yeast strains from numerous sources according to **Table S2**. For the present study, 17 yeast genomes were newly sequenced in our laboratory: DNA samples were processed to generate libraries of short 400-bp inserts. After passing quality control, the libraries were sequenced using an Illumina HiSeq 2000 platform. Sequencing from both ends generated paired-end reads of 2 x 100 bp, resulting in an average sequencing depth of 100X. This dataset was deposited in the European Nucleotide Archive (ENA) under study accessions PRJEB36058. Trimming low quality regions and adapters in Illumina data was performed using Trimmomatic version 0.322 [70] with sequencing parameters: `ILLUMINACLIP:adapterFile:2:30:7`, `LEADING:20`, `TRAILING:20`, `SLIDINGWINDOW:20:25`, `MINLEN:75`.

### Variant calling

We used the Genome Analysis Toolkit (GATK) [71] version 3.6 for SNP and indel calling. Briefly, the workflow is divided into four sequential steps: initial mapping, refinement of the initial reads, multi-sample indel and SNP calling, and finally variant quality score recalibration.

First, reads were aligned to the S288c reference genome (release number R64-1-1, downloaded from SGD) using BWA version 0.7.12 [72] resulting in aligned reads in a BAM file format.

Second, optical and PCR duplicates were removed using MarkDuplicate from the Picard Tools version 2.6.0 (http://picard.sourceforge.net). Base quality scores were recalibrated using BaseRecalibrator/PrintReads (GATK). These recalibrated scores in the output BAM file are closer to their actual probability of mismatching to the reference genome, and are subsequently more accurate. Moreover, the recalibration tool attempts to correct for variation in quality with machine cycle and sequence context. At the end of this step we obtained analysis-ready reads.

Third, we performed SNP and indel discovery using HaplotypeCaller (GATK) on each sample separately in `BP_RESOLUTION` mode, to produce an intermediate file format termed GVCF (for Genomic VCF). These per-sample GVCFs were then run through a joint genotyping step using GenotypeGVCFs (GATK) to produce a raw multi-sample VCF callset.

Fourth, we used Variant Quality Score Recalibration (VQSR) to build an adaptive error model (VariantRecalibrator tool) using an unpublished dataset of known SNPs and Indels obtained from 86 genomes [19]. Then this model was applied (ApplyRecalibration tool) to estimate the probability that each variant in the callset is a true genetic variant or a machine / alignment artifact. This step assigns a VQSLOD score to each variant that is much more reliable than the raw QUAL score calculated by the caller. We used this variant quality score to filter the raw call set, thus producing a subset of calls with our desired level of quality, fine-tuned to balance specificity and sensitivity. This genotyping pipeline resulted in VCF file containing 302,290 biallelic SNPs and 21,045 indels discovered across 68 samples to which were associated a genotyping quality for each strain.

### Population structure

The set 302,290 biallelic SNPs sites identified above was further filtered by removing SNPs with missing genotypes above 0.10, minimum alternate allele frequency (MAF) below 0.03 and SNPs in linkage-disequilibrium using PLINK [73] version 1.9-beta3j with a window size of 50 SNPs, a step of 5 SNPs at a time and a r^2^ threshold of 0.5. The resulting filtered dataset contained 32,379 SNPs positions.

Twenty independent runs of fastStructure version 1.0 [74] were performed varying ancestral population from 1 to 10 using the simple prior. The number of iterations varied from 10 (K = 1) to 70 (K = 10). The highest likelihood was obtained for the solution at 6 ancestral populations (likelihood: -0.75599; total iterations: 30). CLUMPAK [75] was used for analyzing the result of multiple independent runs of fastStructure. CLUMPAK identifies an optimal alignment of inferred clusters across different values of K, simplifying the comparison of clustering results across different K values. Structure plots were obtained using the interactive web application Structure Plot [76].

Structuration was also studied using discriminant analysis of principal components (DAPC) [77] a multivariate method designed to identify clusters of genetically related individuals. This analysis was performed using the R package adegenet version 2.1.1 [78]. DAPC was performed (function dapc) using clusters defined by K-means where we specified a number of clusters from 4 to 7.

### Phylogenetic tree imputation

The set of 302,290 biallelic SNPs sites was filtered by removing SNPs at positions with missing genotypes above 0.10 and minimum alternate allele frequency (MAF) below 0.10. The resulting filtered dataset containd 99,128 SNPs positions. The VCF file was converted into Fasta sequences using `generate_snp_sequence.R` from the R-package SNPhylo [79]. A phylogenetic tree was computed with RAxML version 7.2.8 [80] performing a complete analysis (ML search + 100 bootstrapping) using the `GTRGAMMA` evolution model.

For the microsatellite typing obtained from 229 baker yeasts, Ritland relatedness coefficient [51] was estimated using PolyRelatedness, a software able to estimating pairwise relatedness between individuals with different levels of ploidy [81]. A neighbor joining tree was obtained using the nj function from the ape R package.

### Analysis across the 1,011 genomes data

A genotyping matrix was constructed with the GenotypeGVCFs function of GATK that was run with 1,011 gvcf files constructed in [12] as well as the 26 bakery strains gvcf files that were not included in this previous study (17 newly sequenced diploid sourdough strains genomes and a 9 previously sequenced bakery strains). This extended the dataset to a total of 68 bakery yeast genomes.

The neighbor-joining tree was constructed with the R packages ape and SNPrelate. The gvcf matrix was first converted into a gds file and individual dissimilarities were estimated for each pair of individuals with the snpgdsDiss function. The bionj algorithm was then run on the distance matrix that was obtained.

A set of 552,093 biallelic SNPs was obtained from the gvcf matrix selecting 157 genomes (including 68 bakery strains; **Table S6**). SNPs were further filtered by removing SNPs with missing genotypes above 0.10, minimum alternate allele frequency (MAF) below 0.03 and SNPs in linkage-disequilibrium using PLINK version 1.9-beta3j with a window size of 50 SNPs, a step of 5 SNPs at a time and r^2^ threshold of 0.5. The resulting filtered dataset contains 49,482 SNPs positions. Twenty independent runs of Admixture version 1.3.0 [82] were performed varying ancestral population from 2 to 20. The value of K = 17 exhibited the lowest cross-validation error compared to other K values. CLUMP [83] was used for analysis the results of multiple independent runs of Admixture. Structure plots were obtained using the interactive web application Structure Plot [76].

To evaluate the bakery strains admixture, we also used the TreeMix algorithm [45], which builds population trees and tests for the presence of gene flow between populations. We estimated a maximum-likelihood tree (**Figure S2**) rooted using the China population (CHNV), the likely ancestral population of *S. cerevisiae* [13]. To test for gene flow, we used the three-population f3 test [84] as suggested by [45].

## Acknowledgments

We acknowledge Pierre Gomes for curation of microsatellite data, a panel of bakers and farmer-bakers for providing sourdoughs, F. Minervini and the university of Perugia for providing Italian sourdough strains as well as all the international yeast collections for providing strains. We are also grateful to Agnes Masquin and Isabelle Delmas for their administrative support and Jean-Luc Legras for discussion and critical reading of the manuscript.

## Author contributions

D.Si. supervised the project and designed the experiment; F.B., T.N. and D.Si. analyzed the data and wrote the manuscript. D.Se. performed fitness experiments and conducted fermentation in synthetic medium; T.N. and D.Si. performed statistical analysis on fitness and fermentation data; S.G. was in charge of the yeast collection and genomic DNA extraction for sequencing; L.H., N.A. and A.B. performed microsatellite typing and cytometry analysis; F.B. and A.F. performed genomes analysis.

## Figures

**Figure S1**: Structure-like plot of the probability of membership obtained from 33,032 biallelic SNPs from 68 bakery strains using DAPC. Function dapc was used with clusters defined by K-means where we specified a number of clusters from 4 to 7. The comparison of the final assignments of individuals to groups obtained with fastStructure revealed that the same genetic groups could be recovered.

**Figure S2**: Maximum-likelihood tree of genetic relationships among populations. Branch lengths are proportional to drift in allele frequencies between populations. The scale with the standard error (s.e.) was extracted from the sample covariance matrix. The red arrow shows a migration event resulting in admixture that passed the significance threshold of the three-population test (f3).

**Figure S3**: Relative sequencing depth measured for genes involved in maltose, iso-maltose and sucrose assimilation. Data were obtained for strains selected in the “Mixed origin” clade (49 controls and 35 bakery yeasts) and “Mosaic region 3” clade (79 controls and 18 bakery yeasts). **A**, maltose genes cluster *MAL1-3*; **B**, maltose genes cluster *MAL3-3*; **C**, iso-maltases genes *IMA1-5* and **D**, invertase gene *SUC2*.

**Figure S4:** Population size variation after 27h of fermentation in a sourdough synthetic medium among sourdough, commercial and non-bakery strains (other).

