## Supplemental Fig & Tables for "Evidence for two main domestication trajectories in *Saccharomyces cerevisiae* linked to distinct bread-making processes"

### Figure S1

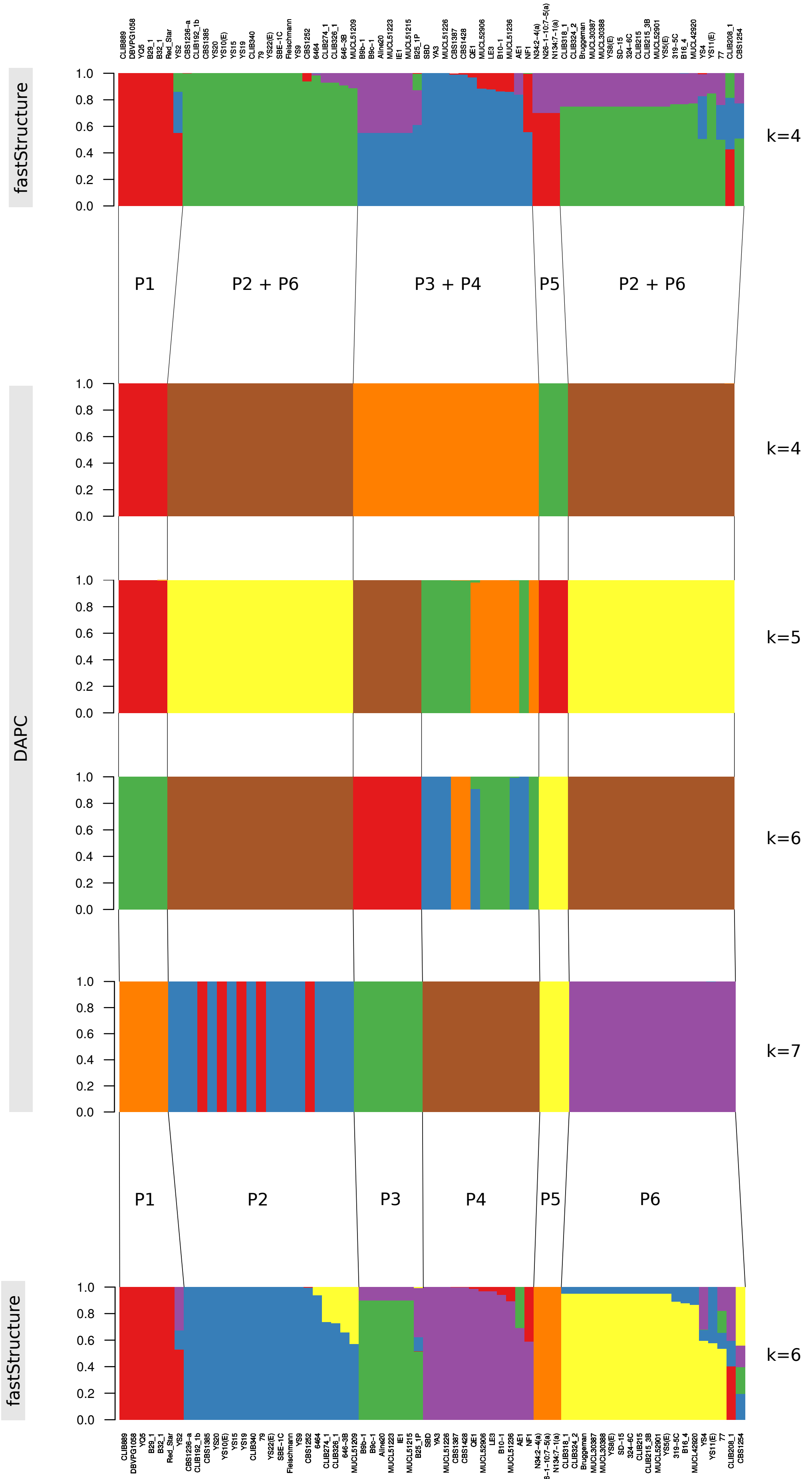
fastStructure
k=4
P1
P2 + P6
P3 + P4
P5
P2 + P6
DAPC
k=4
k=5
k=6
k=7
P1
P2
P3
P4
P5
P6
fastStructure
k=6
1.0
0.8
0.6
0.4
0.2
0.0

### Figure S2

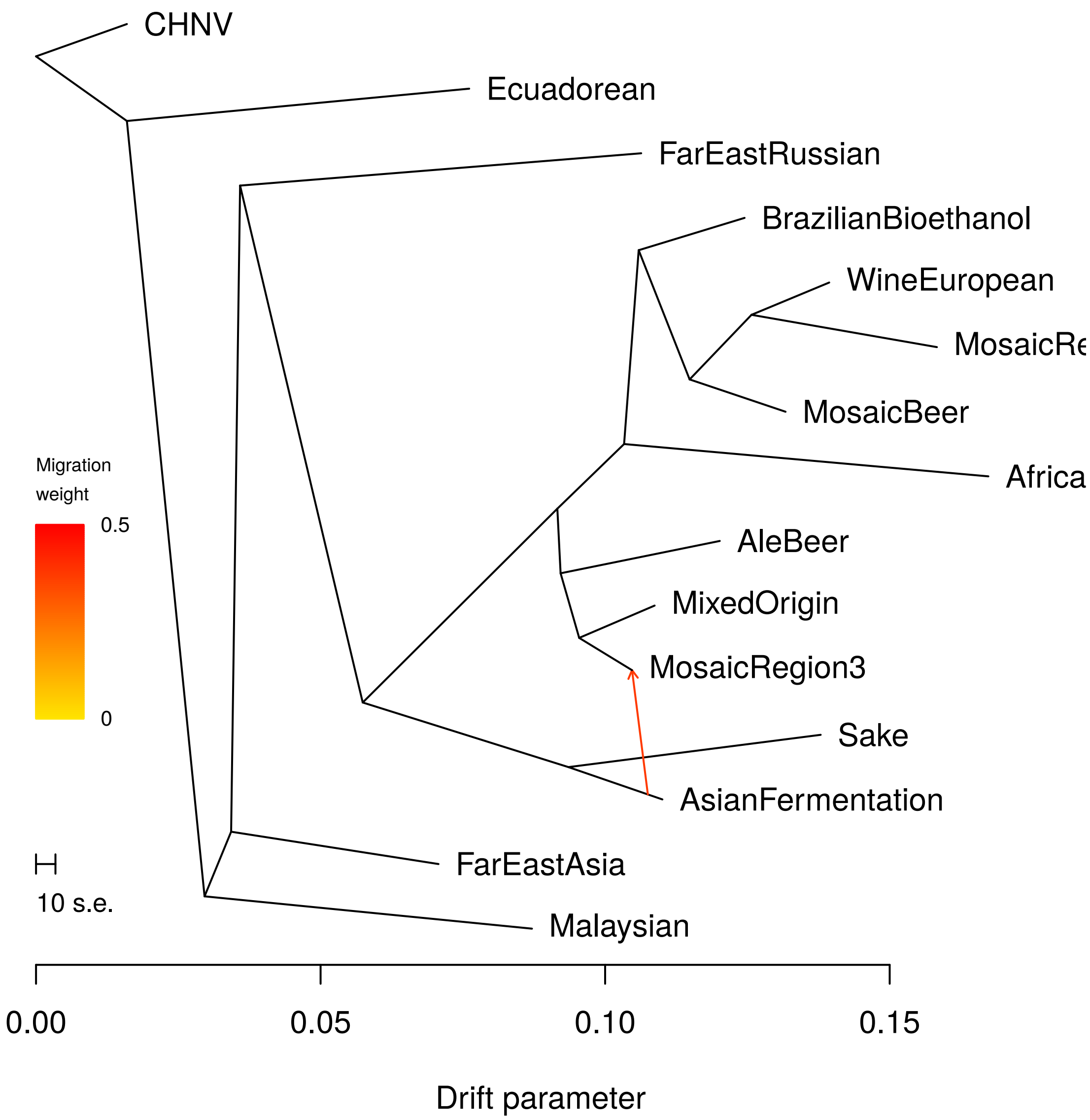

CHNV
Ecuadorean
FarEastRussian
BrazilianBioethanol
WineEuropean
MosaicRe
MosaicBeer
Africa
AleBeer
MixedOrigin
MosaicRegion3
Sake
AsianFermentation
FarEastAsia
Malaysian
Migration weight
0.5
0
10 s.e.
0.00
0.05
0.10
0.15
Drift parameter

Figure S3

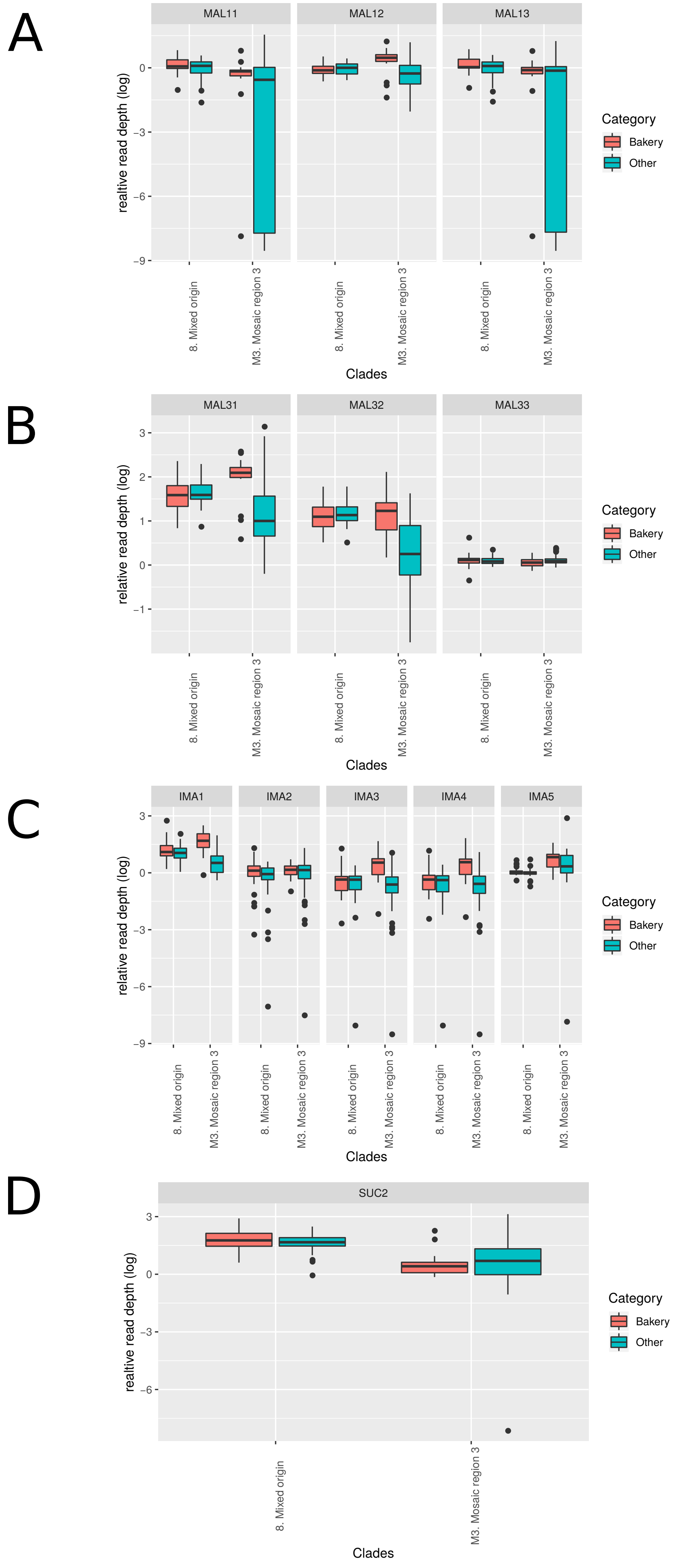

A
MAL11
MAL12
MAL13
realtive read depth (log)
0
−3
−6
−9
8. Mixed origin
M3. Mosaic region 3
Clades
Category
Bakery
Other
B
MAL31
MAL32
MAL33
relative read depth (log)
3
2
1
0
−1
C
IMA1
IMA2
IMA3
IMA4
IMA5
D
SUC2

### Figure S4

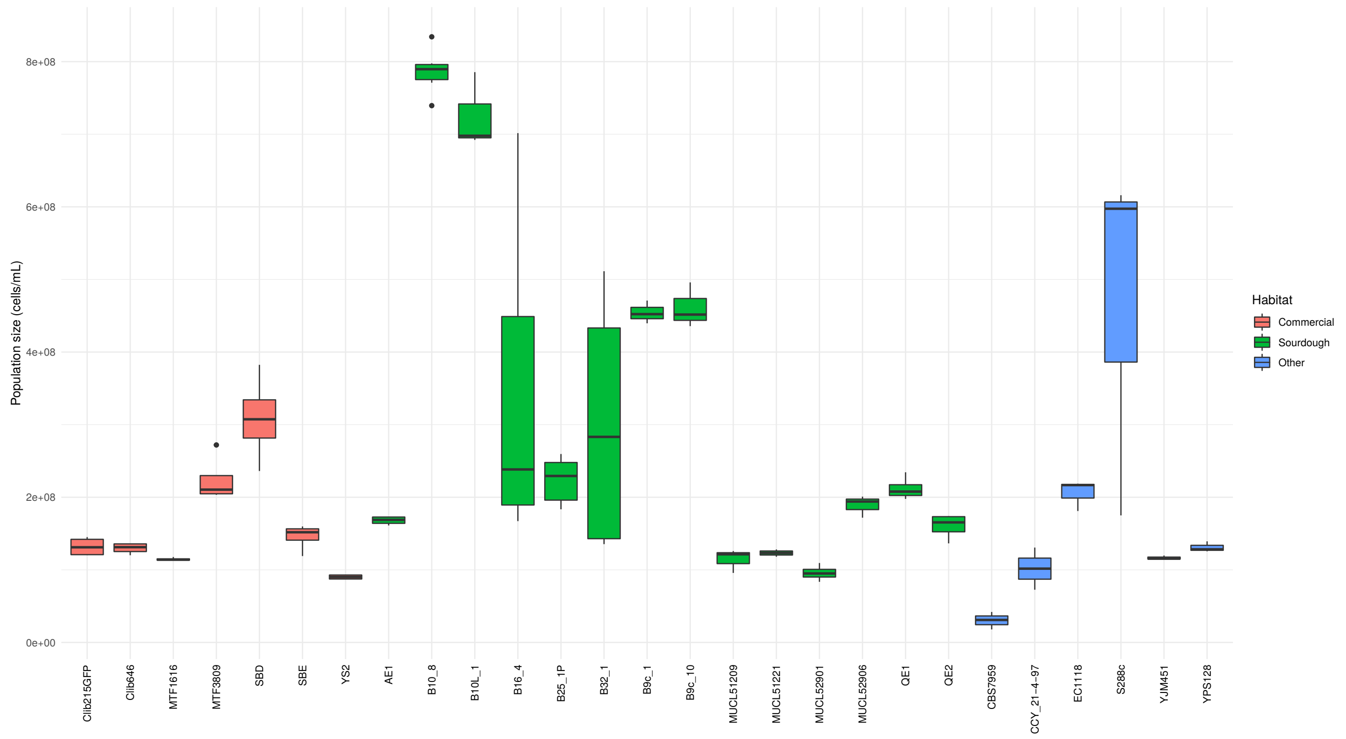

Population size (cells/mL)
0e+00
2e+08
4e+08
6e+08
8e+08
Clib215GFP
Clib646
MTF1616
MTF3809
SBD
SBE
YS2
AE1
B10_8
B10L_1
B16_4
B25_1P
B32_1
B9c_1
B9c_10
MUCL51209
MUCL51221
MUCL52901
MUCL52906
QE1
QE2
CBS7959
CCY_21-4-97
EC1118
S288c
YJM451
YPS128
Habitat
Commercial
Sourdough
Other

TableS2. Origin and accession number of strains used to test for ploidy effect on fitness and fermentation kinetics

| **Strain's name** | **Lab Code** | **Ploidy** | **Competitor(s)** | **Isolation** | **Origin** | **Reference** |
|---|---|---|---|---|---|---|
| 6464 | 2387 | 2n | 6464 GFP | Commercial | Japan | na |
| Clib215 | 2397 | 4n | Clib215 GFP | Commercial | New Zealand | Albertin *et al.* (2009) |
| Clib646 | 3621 | 4n | B32_1 GFP | commercial | USA | Albertin *et al*. (2009) |
| MTF1616 (YS2) | 1616 | 4n | B10L_1 GFP | Commercial | Australia | Liti *et al.* (2009) |
| MTF3809 (Hirondelle) | 3809 | 2n | Clib215 GFP | Commercial | Northern Europe | Albertin *et al.* (2009) |
| MTF4554 (Fleischman) | 4554 | 4n | 6464 GFP | Commercial | USA | Albertin *et al.* (2009) |
| SBD | 3631 | 2n | B16_4 GFP | Commercial | Northern Europe | Albertin *et al.* (2009) |
| SBE | 3629 | 4n | B9b_10 GFP | Commercial | Northern Europe | Albertin *et al.* (2009) |
| 6464 GFP | 4612 | 2n | MTF4554; MUCL52901; 6464 | NA | Laboratory | *This study* |
| B10L_1 GFP | 4611 | 2n | MTF1616; QE2; B10L_1 | NA | Laboratory | *This study* |
| B16_4 GFP | 4608 | 4n | SBD; B25_1P; B16_4 | NA | Laboratory | *This study* |
| B32_1 GFP | 4610 | 4n | QE1; B31_1 | NA | Laboratory | *This study* |
| B9b_10 GFP | 4609 | 2n | MUCL51221; SBE; B9b_10 | NA | Laboratory | *This study* |
| Clib215 GFP | 3280 | 4n | AE1; MTF3809; Clib215 | NA | Laboratory | *This study* |
| AE1 | 3989 | 2n | Clib215 GFP | Sourdough | Italy | Minervini et al. (2012) |
| B10L_1 | 3622 | 2n | B10L_1 GFP | Sourdough | France | Urien *et al.* (2019) |
| B16_4 | 4030 | 4n | B16_4 GFP | Sourdough | France | Michel *et al.* (2019) |
| B25_1P | 3969 | 2n | B16_4 GFP | Sourdough | France | Michel *et al.* (2019) |
| B32_1 | 3972 | 4n | B32_1 GFP | Sourdough | France | Michel *et al.* (2019) |
| B9b_10 | 3976 | 2n | B9b_10 GFP | Sourdough | France | Urien *et al.* (2019) |
| MTF3642 (MUCL 52901 | 3642 | 4n | 6464 GFP | Sourdough | Belgium | Vrancken *et al.* (2010) |
| MUCL51221 | 3624 | 4n | B9b_10 GFP | Sourdough | Belgium | Vrancken *et al.* (2010) |
| QE1 | 3692 | 2n | B32_1 GFP | Sourdough | Italy | Minervini et al. (2012) |
| QE2 | 4619 | 4n | B10L_1 GFP | Sourdough | Italy | Minervini et al. (2012) |

Table S3. Effect of GFP labeling on relative fitness (w)

| **Souche_GFP** | **t** | **df** | **P-value** | **Mean** |
|---|---|---|---|---|
| CLIB_215_GFP | -2,1301 | 2 | 0,1669 | 0,9817 |
| B16_4_GFP | 2,7451 | 2 | 0,1110 | 1,0058 |
| B9_10_GFP | 0,8098 | 1 | 0,5667 | 1,0059 |
| B32_1_GFP | 0,1994 | 2 | 0,8604 | 1,0012 |
| B10_1_GFP | 0,8483 | 2 | 0,4856 | 1,0028 |
| 6464_GFP | 5,9195 | 2 | 0,0274 | 1,0251 |

Table S4. Effect of GFP labelling on fermentation kinetics

| Strain | Term | df | Sum sq | Mean sq | F | P-value | Parameter |
|---|---|---|---|---|---|---|---|
| B32_1 | GFP | 1 | 0,1873 | 0,1873 | 0,1567 | 0,7124 | t_1g |
| | Residuals | 4 | 4,7790 | 1,1948 | | | |
| B32_1 | GFP | 1 | 0,2400 | 0,2400 | 1,3059 | 0,3169 | CO2max |
| | Residuals | 4 | 0,7351 | 0,1838 | | | |
| B32_1 | GFP | 1 | 0,0283 | 0,0283 | 0,1020 | 0,7655 | Vmax |
| | Residuals | 4 | 1,1097 | 0,2774 | | | |
| B32_1 | GFP | 1 | 0,2509 | 0,2509 | 0,0799 | 0,7914 | tVmax |
| | Residuals | 4 | 12,5598 | 3,1400 | | | |
| B10_1 | GFP | 1 | 1,6527 | 1,6527 | 0,6683 | 0,4595 | t_1g |
| | Residuals | 4 | 9,8922 | 2,4731 | | | |
| B10_1 | GFP | 1 | 0,0131 | 0,0131 | 0,0205 | 0,8930 | CO2max |
| | Residuals | 4 | 2,5439 | 0,6360 | | | |
| B10_1 | GFP | 1 | 0,1014 | 0,1014 | 8,0222 | 0,0472 | Vmax |
| | Residuals | 4 | 0,0506 | 0,0126 | | | |
| B10_1 | GFP | 1 | 4,7615 | 4,7615 | 1,3233 | 0,3141 | tVmax |
| | Residuals | 4 | 14,3925 | 3,5981 | | | |
| B16_4 | GFP | 1 | 2,9906 | 2,9906 | 0,6632 | 0,4611 | t_1g |
| | Residuals | 4 | 18,0378 | 4,5094 | | | |
| B16_4 | GFP | 1 | 0,0000 | 0,0000 | 0,0001 | 0,9942 | CO2max |
| | Residuals | 4 | 1,0959 | 0,2740 | | | |
| B16_4 | GFP | 1 | 0,1803 | 0,1803 | 2,5783 | 0,1836 | Vmax |
| | Residuals | 4 | 0,2797 | 0,0699 | | | |
| B16_4 | GFP | 1 | 7,4884 | 7,4884 | 2,1374 | 0,2176 | tVmax |
| | Residuals | 4 | 14,0140 | 3,5035 | | | |
| B9_10 | GFP | 1 | 1,6443 | 1,6443 | 0,4661 | 0,5323 | t_1g |
| | Residuals | 4 | 14,1109 | 3,5277 | | | |
| B9_10 | GFP | 1 | 0,1441 | 0,1441 | 0,2936 | 0,6167 | CO2max |
| | Residuals | 4 | 1,9637 | 0,4909 | | | |
| B9_10 | GFP | 1 | 0,0001 | 0,0001 | 0,0190 | 0,8969 | Vmax |
| | Residuals | 4 | 0,0219 | 0,0055 | | | |
| B9_10 | GFP | 1 | 0,1663 | 0,1663 | 0,1354 | 0,7315 | tVmax |
| | Residuals | 4 | 4,9122 | 1,2281 | | | |
| 6464 | GFP | 1 | 2,5117 | 2,5117 | 0,8594 | 0,4064 | t_1g |
| | Residuals | 4 | 11,6899 | 2,9225 | | | |
| 6464 | GFP | 1 | 0,5828 | 0,5828 | 6,9994 | 0,0572 | CO2max |
| | Residuals | 4 | 0,3331 | 0,0833 | | | |
| 6464 | GFP | 1 | 0,0610 | 0,0610 | 1,6255 | 0,2713 | Vmax |
| | Residuals | 4 | 0,1501 | 0,0375 | | | |
| 6464 | GFP | 1 | 9,5407 | 9,5407 | 4,9699 | 0,0897 | tVmax |
| | Residuals | 4 | 7,6789 | 1,9197 | | | |
| CLIB_215 | GFP | 1 | 0,0243 | 0,0243 | 0,1862 | 0,6883 | t_1g |
| | Residuals | 4 | 0,5223 | 0,1306 | | | |
| CLIB_215 | GFP | 1 | 0,2400 | 0,2400 | 0,8033 | 0,4208 | CO2max |
| | Residuals | 4 | 1,1950 | 0,2988 | | | |
| CLIB_215 | GFP | 1 | 0,0010 | 0,0010 | 0,0141 | 0,9113 | Vmax |
| | Residuals | 4 | 0,2883 | 0,0721 | | | |
| CLIB_215 | GFP | 1 | 0,0466 | 0,0466 | 1,3322 | 0,3127 | tVmax |
| | Residuals | 4 | 0,1400 | 0,0350 | | | |

### Table S7. Gross chromosomal rearrangements or aneuploidies

| ID | Name | Origin | Clades | Type | chr01 | chr02 | chr03 | chr04 | chr05 | chr06 | chr07 | chr08 | chr09 | chr10 | chr11 | chr12 | chr13 | chr14 | chr15 | chr16 |
|---|---|---|---|---|---|---|---|---|---|---|---|---|---|---|---|---|---|---|---|---|
| BTF | YS22(E) | Bakery | 8. Mixed origin | Commercial | x | | | | | | | | x | | | | | | | |
| CBF | SD-15 | Bakery | 8. Mixed origin | Sourdough | | | | | | | | | x | | | | | | | |
| CET | N134:7-1(a) | Bakery | 6. African beer | Dough | | | | | | | | | x | | | | | | | |
| ARB | CBS1254 | Bakery | M3. Mosaic region 3 | Commercial | | | | | | | | x | | x | | | | | | |
| ASL | CLIB340 | Bakery | 8. Mixed origin | Sourdough | | x | | | | | | | x | | | | | | | |
| CHH | 319-5C | Bakery | 8. Mixed origin | Commercial | | | | | | | x | | | | R | | | | | |
| APN | YS19 | Bakery | 8. Mixed origin | Commercial | x | | | | | | | | | | | | | | | |
| BTB | 77 | Bakery | M3. Mosaic region 3 | Commercial | x | | | | R | | | x | | | L | | | | | |
| ADK | CLIB208_1 | Bakery | M3. Mosaic region 3 | NA | | | | | | | | | x | | | | | | | |
| ADL | CLIB318_1 | Bakery | 8. Mixed origin | Commercial | x | | | | | x | | | x | | x | | | | | |
| AGE | CBS1385 | Bakery | 8. Mixed origin | Commercial | | | | | | x | | | | | | | | | | |
| API | YS20 | Bakery | 8. Mixed origin | Commercial | | | | | | | | x | x | | | | | | | |
| APM | YS15 | Bakery | 8. Mixed origin | Commercial | | | | | | | | | x | | | | | | | |
| APL | YS11(E) | Bakery | 8. Mixed origin | Commercial | | | | | | | L | | | | | | | | | |
| CHI | 646-3B | Bakery | 8. Mixed origin | Commercial | | | R | | | | | | | | | | | | | |
| CHL | SBE-1C | Bakery | 8. Mixed origin | Commercial | | x | | | | | | | | | | | x | | | |
| AAL | CBS1236-a | Bakery | 8. Mixed origin | Commercial | | | | | | x | | | | | | | | | | |
| AAP | CLIB326_1 | Bakery | 8. Mixed origin | Commercial | | | | | | | | | | | | | | R | | |
| CES | N26-1-10:7-5(a) | Bakery | 6. African beer | Dough | x | | | | | | | | | | | | | | | |
| AAN | CLIB274_1 | Bakery | 8. Mixed origin | Commercial | | | x | | | | x | | | | | | | | | |
| SRR3265414 | Red Star | Bakery | 1. Wine/European | Commercial | | x | | | x | | | | x | | | | | | | |
| BR004 | MUCL42920 | Bakery | 8. Mixed origin | Commercial | | | | | | | | | x | | x | | | | | |
| MTF3695 | IE1 | Bakery | M3. Mosaic region 3 | Sourdough | | | x | | | | | | | | | | | | | |
| MTF3690 | MUCL51226 | Bakery | 26. Asian fermentation | Sourdough | R | | | | | | | | | | | | | | | |
| MTF3632 | Aline20 | Bakery | M3. Mosaic region 3 | Sourdough | x | | | | | | | | | | | | | | | |
| PYg78 | MUCL52901 | Bakery | 8. Mixed origin | Sourdough | x | | | | | | | | | | | | | | | |
| | | | | Numbers | 8 | 3 | 3 | 0 | 2 | 3 | 3 | 3 | 10 | 1 | 4 | 0 | 1 | 1 | 0 | 0 |

TableS8. Origin, accession number and genomic clades of phenotyped strains

| Strain's name | Isolation | Lab Code | Origin | Genomic Sub_Gro | Genomic Group | Provider-Associated paper |
|---|---|---|---|---|---|---|
| YPS 128 | American oak | / | USA | NA | North American oak | Liti et al. (2009), Peter et al. 2019 |
| CBS 7959 | Bioethanol | 2351 | Brazil | NA | Brazilian bioethanol | Liti et al. (2009), Peter et al. 2019 |
| YJM 451 | Clinical | 1785 | Europe | NA | Mosaic region 3 | Liti et al. (2009), Peter et al. 2019 |
| EC1118 | Winery | 1309 | France | NA | Wine | Commercial |
| CCY_21-4-97 | Fish pond water | 2389 | Slovakia | NA | Mixed origin | Liti et al. (2009), Peter et al. 2019 |
| S288c | Rotting fig - Lab | 1599 | California | NA | NA | Liti et al. (2009), Peter et al. 2019 |
| 6464 | Commercial | 2387 | Japan | P2_green | Mixed origin | Commercial |
| MTF3809 | Commercial | 3809 | Northern Eu | NA | NA | Commercial |
| YS2 | Commercial | 1616 | Australia | P1_red | Mosaic region 3 | Liti et al. (2009), Peter et al. 2019 |
| Clib646 | Commercial | 3621 | USA | P2_green | Mixed origin | CIRM-Levure |
| SBE | Commercial | 3629 | Northern Eu | P2_green | Mixed origin | Commercial-Albertin et al. 2009 |
| MTF1616 | Commercial | 1616 | Australia | P2_green | Mixed origin | Commercial |
| SBD | Commercial | 3631 | Northern Eu | P4_blue | Mosaic region 3 | Commercial-Albertin et al. 2009 |
| Clib215 | Commercial | 2397 | New Zealand | P6_purple | Mixed origin | CIRM-Levure |
| MUCL51209 | Sourdough | 3992 | Belgium | P2_green | Mixed origin | MUCL collection -Vrancken et al. 2009 |
| MUCL52906 | Sourdough | 3995 | Belgium | P4_blue | Mosaic region 3 | MUCL collection -Vrancken et al. 2009 |
| MUCL52901 | Sourdough | 3689 | Belgium | P6_purple | Mixed origin | MUCL collection -Vrancken et al. 2009 |
| MUCL51221 | Sourdough | 3624 | Belgium | NA | NA | MUCL collection -Vrancken et al. 2009 |
| B32_1 | Sourdough | 3972 | France | P1_red | Mosaic region 1 | Baker B33-Michel et al. 2019 |
| B25_1P | Sourdough | 3969 | France | P3_yellow | Mosaic region 3 | Baker B25-Michel et al. 2019 |
| B9c_1 | Sourdough | 4575 | France | P3_yellow | Mosaic region 3 | Baker B9-Urien et al. 2018 |
| B9c_10 | Sourdough | 3976 | France | P3_yellow | Mosaic region 3 | Baker B9-Urien et al. 2018 |
| B10_8 | Sourdough | 4599 | France | P4_blue | Mosaic region 3 | Baker B10-Urien et al 2018 |
| B10L_1 | Sourdough | 3622 | France | P4_blue | Mosaic region 3 | Baker B10-Urien et al 2018 |
| B16_4 | Sourdough | 4030 | France | P6_purple | Mixed origin | Baker B16-Michel et al 2019 |
| AE1 | Sourdough | 3989 | Italy | P4_blue | Mosaic region 3 | Perugia collection-Minervini et al. 2012 |
| QE1 | Sourdough | 3692 | Italy | P4_blue | Mosaic region 3 | Perugia collection-Minervini et al. 2012 |
| QE2 | Sourdough | 4619 | Italy | NA | NA | Perugia collection-Minervini et al. 2012 |

Table S9. Statistical tests of the "habitat of origin" (commercial/sourdough/other) effect on fermentation parameters and on the population size after 27h of fermentation.

| t1g | Sum sq | Mean sq | NumF | DenDF | F value | Pr (>F) |
|---|---|---|---|---|---|---|
| Habitat | 23.66 | 11.83 | 2 | 27.67 | 3.6838 | 0.038 |
| | Mean | SE | Tukey group | | | |
| Commercial | 6.17 | 1.82 | c | | | |
| Sourdough | 8.64 | 2.49 | b | | | |
| Other | 11.28 | 2.15 | a | | | |
| CO2max | Sum sq | Mean sq | NumF | DenDF | F value | Pr (>F) |
| Habitat | 2040.2 | 1020.1 | 2 | 95.26 | 50.28 | 1.2. 10-15 |
| | Mean | SE | Tukey | | | |
| Commercial | 27.51 | 5.03 | a | | | |
| Sourdough | 24.42 | 4.31 | b | | | |
| Other | 10.04 | 5.79 | c | | | |
| Vmax | Sum sq | Mean sq | NumF | DenDF | F value | Pr (>F) |
| Habitat | 10.374 | 5.1868 | 2 | 23.684 | 37.1 | 5.04 10-8 |
| | Mean | SE | | | | |
| Commercial | 2.87 | 0.36 | a | | | |
| Sourdough | 2.68 | 0.68 | a | | | |
| Other | 1.16 | 0.52 | b | | | |
| tVmax | Sum sq | Mean sq | NumF | DenDF | F value | Pr (>F) |
| Habitat | 3.5961 | 1.7981 | 2 | 27.254 | 0.2709 | 0.7647 |
| Commercial | 20.04 | 3.69 | | | | |
| Sourdough | 19.28 | 3.95 | | | | |
| Other | 17.78 | 4.00 | | | | |
| Cellt27 | Sum sq | Mean sq | NumF | DenDF | F value | Pr (>F) |
| Habitat | 4.3887 | 2.1943 | 2 | 27866 | 9.9215 | 4.93 10-5 |
| | Mean | SE | | | | |
| Commercial | 1.63. 10^8 | 0.7. 10^8 | b | | | |
| Sourdough | 3.18. 10^8 | 2.4. 10^8 | a | | | |
| Other | 1.79. 10^8 | 1.6 10^8 | b | | | |

TableS10. Amova results

```
==============================================================
AMOVA (structuration: commercial vs sourdough strains)
==============================================================

poppr.amova(total, hier = ~Isolation, cutoff = .20, freq = TRUE)

$results
                 Df     Sum Sq   Mean Sq
Between samples   1   37.60309 37.603091
Within samples  227 1086.09778  4.784572
Total           228 1123.70087  4.928513

$componentsofcovariance
                               Sigma         %
Variations  Between samples 0.612206  11.34392
Variations  Within samples  4.784572  88.65608
Total variations            5.396778 100.00000

$statphi
                        Phi
Phi-samples-total 0.1134392

Monte-Carlo test
Call: as.randtest(sim = res, obs = sigma[1])

Observation: 0.612206

Based on 1000 replicates
Simulated p-value: 0.000999001
Alternative hypothesis: greater

     Std.Obs  Expectation     Variance
14.288937405 -0.001856591  0.001846824

==============================================================
AMOVA (structuration by country, only 3 strains by sourdough )
==============================================================

poppr.amova(sampled.sourdough.europe, hier = ~Country, cutoff = .20,
freq = TRUE)

$results
                Df    Sum Sq   Mean Sq
Between samples  2  40.33112 20.165562
Within samples  90 450.88393  5.009821
Total           92 491.21505  5.339294

$componentsofcovariance
                                Sigma          %
Variations  Between samples 0.5255346   9.494143
Variations  Within samples  5.0098214  90.505857
```

```
Total variations            5.5353561 100.000000

$statphi
                          Phi
Phi-samples-total 0.09494143

Monte-Carlo test
Call: as.randtest(sim = res, obs = sigma[1])

Observation: 0.5255346

Based on 1000 replicates
Simulated p-value: 0.000999001
Alternative hypothesis: greater

     Std.Obs  Expectation     Variance
10.580045810  0.001102749  0.002456988
```

```
=======================================================
AMOVA (structuration by country, all sourdough strains)
=======================================================

poppr.amova(sourdough.europe, hier = ~Country, cutoff = .20, freq =
TRUE)

$results
                 Df    Sum Sq   Mean Sq
Between samples   2  95.23948 47.619738
Within samples  195 900.60427  4.618483
Total           197 995.84375  5.055044

$componentsofcovariance
                                Sigma        %
Variations  Between samples 0.8616788  15.7236
Variations  Within samples  4.6184835  84.2764
Total variations            5.4801623 100.0000

$statphi
                        Phi
Phi-samples-total 0.157236

Monte-Carlo test
Call: as.randtest(sim = res, obs = sigma[1])

Observation: 0.8616788

Based on 1000 replicates
Simulated p-value: 0.000999001
Alternative hypothesis: greater

      Std.Obs   Expectation      Variance
25.8415416163 -0.0002726502  0.0011125737
```

```
=================================================================
AMOVA (structuration by sourdough, only french sourdough strains)
=================================================================

poppr.amova(sourdough.france, hier = ~Sourdough, cutoff = .20, freq
= TRUE)

$results
                 Df   Sum Sq   Mean Sq
Between samples  11 410.7423 37.340205
Within samples  117 155.1628  1.326178
Total           128 565.9050  4.421133

$componentsofcovariance
                               Sigma         %
Variations  Between samples 3.669149  73.45163
Variations  Within samples  1.326178  26.54837
Total variations            4.995326 100.00000

$statphi
                        Phi
Phi-samples-total 0.7345163

Monte-Carlo test
Call: as.randtest(sim = res, obs = sigma[1])

Observation: 3.669149

Based on 1000 replicates
Simulated p-value: 0.000999001
Alternative hypothesis: greater

      Std.Obs   Expectation      Variance
48.1595021343 -0.0002342938  0.0058052664
```
